# Pharmacological intervention targeting neuroimmune axis in the aging hypothalamus prevents age-associated physiological decline

**DOI:** 10.64898/2026.01.12.698951

**Authors:** Min-Jong Seok, Na-Kyung Lee, In-Seop Sim, Jumi Kim, Kelvin Pieknell, Thor D Stein, Jung-Hee Lee, Hoon Ryu, Seung-Hyup Song, Je-Kyung Seong, Il-Yong Kim, Eun Ha Ko, Yanuar Alan Sulistio, Sang-Hun Lee

## Abstract

Systemic physiological aging is largely driven by disrupted metabolic homeostasis, yet the central mechanisms of this metabolic dysfunction remain poorly defined. Here, we identify the hypothalamus as a critical hub driving systemic aging through neuroimmune-mediated mechanisms. Single-cell transcriptomic and immunohistochemical analyses revealed that aged hypothalami exhibit significant infiltration of CD8⁺ T lymphocytes, beginning in middle age, with signatures of activation and tissue residency. These T cells intimately interact with tanycytes and microglia, promoting neuroinflammation and progressive tanycyte loss, a defining hallmark of hypothalamic aging. T cell receptor profiling revealed a substantial presence of invariant natural killer T (iNKT) and mucosal-associated invariant T (MAIT) cells, likely activated through cytokine-driven, antigen-independent mechanisms. In aged hypothalamus, microglia secrete chemokines CCL3 and CCL4, whose ectopic expression in young mice was sufficient to trigger persistent hypothalamic T cell infiltration and accelerated systemic aging. Circulating oxidized LDLs (OxLDLs) were identified as upstream inducers of this chemokine response. Notably, pharmacological blockade of the CCL3/4-CCR5 axis with Maraviroc and Cenicriviroc prevented T cell recruitment and ameliorated metabolic and physiological impairments. Given the clinical safety of CCR5 antagonists and individuals lacking functional CCR5 remain generally healthy throughout life, our findings highlight midlife CCL3/4-CCR5 inhibition as a translatable therapeutic target for delaying age-related decline and promoting healthspan.

## Introduction

Aging is characterized by progressive physiological decline underpinned by systemic metabolic dysregulation and loss of homeostatic balance. In recent years, extending healthspan, the period of life free from disability, has become a global priority. The United Nations has designated the “Decade of Healthy Aging” as part of its action plan, underscoring the urgency of understanding the biology of aging and developing interventions that mitigate its deleterious effects.

The hypothalamus, located around the third ventricle, serves as a central control hub for whole-body metabolism and homeostasis. By regulating hormone output and modulating the autonomic nervous system, it exerts wide-ranging influence on peripheral organs such as the liver, adipose tissue, skeletal muscle, bone, and intestine. With aging, hypothalamic function deteriorates, disrupting endocrine, autonomic, and neural regulation, and contributing to physiological decline^1–6^. Consequently, the hypothalamus has emerged as a promising target for anti-aging interventions^7^, underscoring the importance of understanding its normal physiology and age-related alterations.

The hypothalamus integrates peripheral metabolic cues through fenestrated capillaries and processes them via neuronal–glial networks. Glial cells, including astrocytes, microglia, and tanycytes, play crucial roles in sensing, relaying, and modulating nutritional and hormonal signals^6,8–10^. With aging, these cells adopt pro-inflammatory states that fuel chronic neuroinflammation. Of particular importance are tanycytes, a neural stem-like population lining the third ventricle. Tanycytes generate neurons^11,12^, regulate orexigenic/anorexigenic circuits^13^, control the hypothalamic–pituitary–thyroid axis^14^, and transmit peripheral metabolic signals^15,16^. Their numbers decline with age^6^, and recent transcriptomic analyses identified tanycytes as among the most transcriptionally altered hypothalamic cells during brain aging^17^. These observations underscore the importance of understanding how hypothalamic glia change with age to advance healthspan-focused strategies.

Emerging work has identified T cell infiltration as a feature of brain aging and age-associated brain disorders. CD8^+^ T cells (CD8) accumulate around the lateral ventricles and white matter, impairing neural stem cells and oligodendrocytes^18,19^. T cell infiltration is also a hallmark of neurodegenerative diseases, including clonal CD8 expansion in Alzheimer’s disease^20,21^ and pathogenic CD4^+^ T cells (CD4) responses in Parkinson’s disease and Lewy body dementia^22^. Yet, whether similar infiltration occurs in the hypothalamus—and whether it contributes to the breakdown of systemic homeostasis during aging—remains unknown.

In this study, we investigated age-dependent changes in the hypothalamus, their contribution to systemic decline, and whether this process is therapeutically targetable. We show here that hypothalamic aging is driven by CD8 orchestrated by senescent microglia through the CCL3/4–CCR5 axis, leading to neuroimmune dysfunction and systemic decline. Targeting this pathway emerges as a potential strategy to preserve hypothalamic integrity and extend healthspan.

## Results

### Age-Dependent Transcriptomic Changes in the Hypothalamus

To characterize the molecular changes that occur during hypothalamic aging, we conducted genome-wide transcriptomic analyses on hypothalamic tissues from mice aged 2, 12, and 26 months (**Fig. 1a**). This time series approach allowed us to identify both early and late transcriptomic events associated with aging. The majority of age-dependent DEGs showed upregulation at 12 and 26 months, with downregulated genes comprising only a small subset. (**Fig. 1b and Extended Data Fig. 1a**). Notably, many DEGs displayed progressively increased expression at 12 and 26 months compared to young controls **(Fig. 1c)**, suggesting a stepwise activation of aging-associated pathways. Most DEGs upregulated at 12 months (27 of 32) remained elevated at 26 months, while 126 additional DEGs emerged uniquely at 26 months **(Extended Data Fig. 1b)**. These findings point to two distinct classes of upregulated DEGs: those activated early and sustained during aging, and those induced only in advanced age, each potentially contributing to different physiological outcomes.

**Fig. 1.**
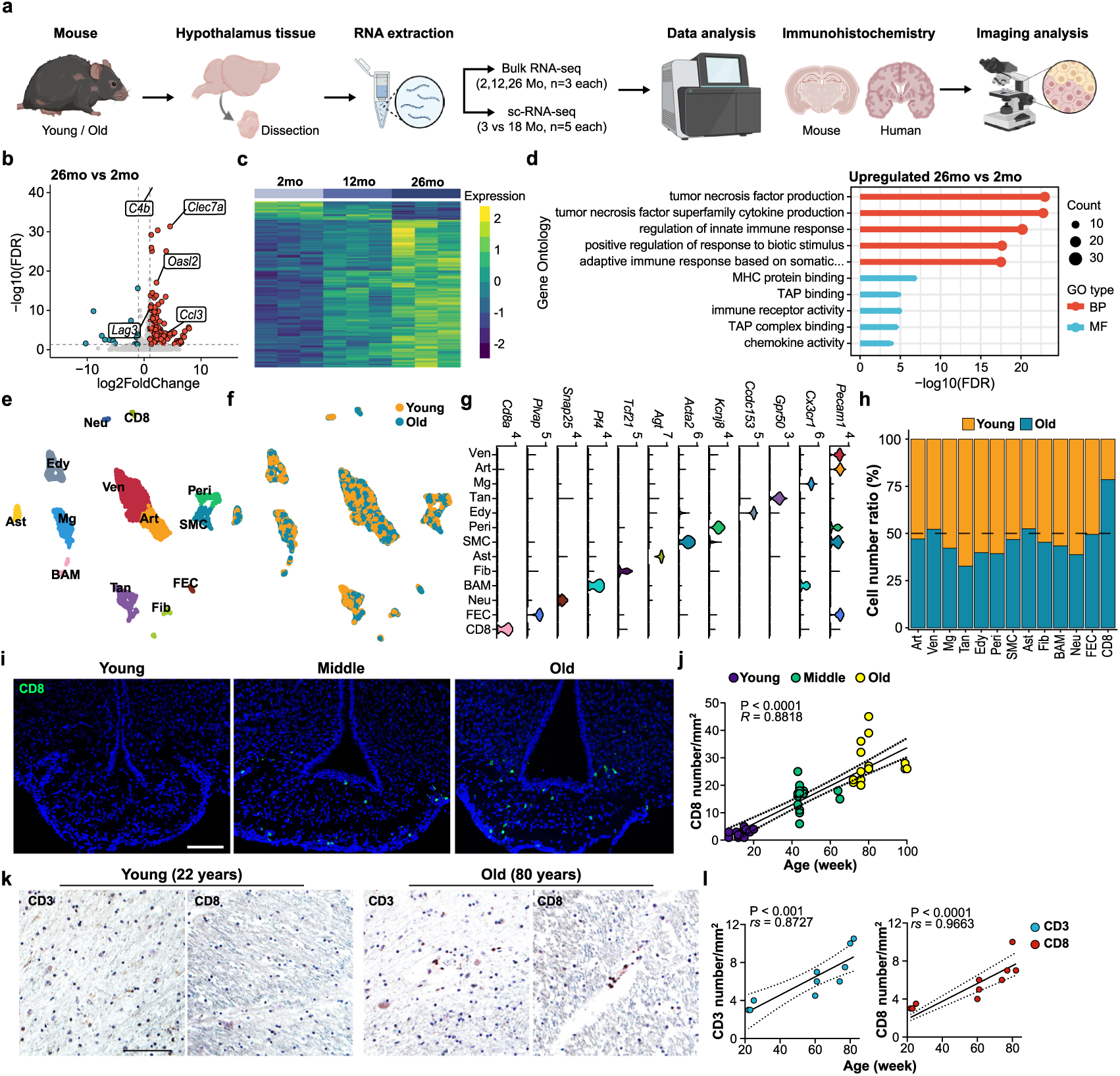
Transcriptomic profiling reveals age-dependent CD8+ T cell infiltration and immune activation in the hypothalamus. **a,** Experimental workflow illustrating dataset generation for Fig. 1. **b-d,** Bulk RNA-seq analysis of hypothalamic tissue from young (2 mo), middle-aged (12 mo), and old (26 mo) mice *(n* = 3 per group). **b,** Volcano plot of differentially expressed genes (DEGs; adjusted *P* < 0.01 and log2FC > 2 or < -2) in 26-mo vs 2-mo mice, with representative immune-related genes highlighted. **c,** Heatmap ofDEGs across all age groups. **d,** Top enriched Gene Ontology (GO) biological process (BP) and molecular function (MF) terms among upregulated genes. **e-h,** Single-cell RNA-seq of hypothalamus from young (3 mo) and old (20 mo) mice *(n* = 5 per group). **e,** UMAP projection of 13 transcriptionally defined cell types: arterial endothelial cell (Art), venous endothelial cell (Ven), microglia (Mg), tanycyte (Tan), ependymal cell (Edy), pericyte (Peri), smooth muscle cell (SMC), astrocyte (Ast), fibroblast (Fib), border-associated macrophage (BAM), neuron (Neu), fenestrated endothelial cell (FEC), and CD8^+^ T cell (CD8). **f,** UMAP projection of young and old hypothalamic cell clusters, showing no overt batch effect. **g,** Violin plots of representative marker genes for each cell cluster. **h,** Proportional changes in each cell type with aging. **i,** Representative immunohistochemical staining of CD8⁺ cells in the hypothalamus of young (7–20 weeks), middle-aged (43–65 weeks), and old (76–100 weeks) mice (*n* = 14, 13, and 16 per group). Scale bar, 100 μm. **j,** Pearson correlation between hypothalamic CD8⁺ cell density (cells/mm²) and age in mice. **k,** Representative immunohistochemical staining of CD3⁺ and CD8⁺ T cells (brown) in postmortem human hypothalamus from young (22 years) and old (80 years) individuals. Scale bar, 100 μm. **l,** Spearman correlation between age and hypothalamic CD3⁺ (left) or CD8⁺ (right) T cell density in human hypothalamus sections (cells/mm²). Data from three young (22–25 years) and seven old (60–82 years) individuals.

Gene Ontology (GO) analysis comparing gene expression between 2 and 26 months identified significant enrichment of immune-related processes, particularly inflammation and adaptive immune responses (**Fig. 1b, d** and **Extended Fig. 1c-f**). Confirming these findings, immunoblot analysis revealed age-dependent elevation of the inflammatory markers IL-1β and TNF-α, together with increased expression of Pan-T cell marker CD3e in aged hypothalamus (**Extended Data Fig. 3a-b**). Consistent with the presence of DEGs activated at both early and late stages of aging (**Extended Data Fig. 1b**), a closer examination of immune-related signatures revealed distinct molecular programs characteristic of different aging phases. At the intermediate aging stage (12 months), there was prominent expression of genes associated with the complement cascade pathway (*C1qa, C3, C4a, C4b*) and Serpina-family protease inhibitors (*Serpina3c, Serpina3n*) (**Extended Data Fig. 1c**), potentially reflecting enhanced debris clearance. This phase also showed elevated expression of MHC class I molecules (*H2-K1, H2-K2, H2-Q4, H2-Q7*) (**Extended Data Fig. 1d**). By the late aging stage (26 months), the immune profile shifted toward increased expression of interferon-stimulated genes (ISGs) (*Ifi207, Ifi44, Isg15, Oas1a, Oas2, Stat1*), indicating a transition toward adaptive immune responses and potential T-cell infiltration (*Lag3, Lilrb4a, Mmp12, Lyz2, Ly9*) (**Extended Data Fig. 1e-f**). This biphasic aging pattern aligns closely with human biomarker studies^23^. Specifically, a previous study identified two peaks (referred to as Crest 1 and Crest 2) in human aging: Crest 1, occurring around 44 years of age (∼12 months in mice), featured prominent activation of the complement cascade pathway and MHC class I expression, a finding that persisted into Crest 2 (approximately 60 years in humans, corresponding to ∼20–24 months in mice). However, interferon-response genes, such as *Stat1*, emerged specifically in the later aging stage (Crest 2), further supporting the distinct transition as we observed in our bulk RNA-seq data.

### CD8 infiltration defines the aged hypothalamus

To better understand the cellular heterogeneity underlying these transcriptomic changes, we performed single-cell RNA sequencing (scRNA-seq) on pooled hypothalamic tissues from mice aged 3 and 18 months. After quality control and removal of doublets (**Extended Data Fig. 2a**), 8,184 cells remained for downstream analysis and were visualized using Uniform Manifold Approximation and Projection (UMAP) (**Fig. 1e).** Unsupervised clustering identified 13 distinct cellular populations, manually annotated based on established marker genes and validated by automated sc-type classification (**Supplementary Fig. 1)**. These populations included venous endothelial cells (Ven), arterial endothelial cells (Art), microglia (Mg), tanycytes (Tan), ependymal cells (Edy), pericytes (Peri), vascular smooth muscle cells (SMC), astrocytes (Ast), fibroblasts (Fib), border-associated macrophages (BAM), neurons (Neu), fenestrated endothelial cells (FEC), and CD8⁺ T cells (CD8) (**Fig. 1e, g and Extended Data Fig. 2b-d**). Data integration confirmed no detectable batch effects (**Fig. 1f and Extended Data Fig. 2a**). GO enrichment analysis of cluster-specific marker genes supported the accuracy of cell-type annotations (**Supplementary Fig. 2**). The most striking population shift in scRNA-seq data was a robust increase of CD8 cluster in the aged hypothalamus (**Fig. 1h**). This aligns with the bulk RNA-seq findings that identified GO associated with ‘‘adaptive immune response based on somatic recombination of immune receptors built from immunoglobulin superfamily domains”, “MHC protein binding” as the most prominently enriched pathways in aging (**Fig. 1d**).

Immunohistochemical analysis confirmed preferential accumulation of CD8 but notably not CD4 in the hypothalamic parenchyma of aged mice (**Fig. 1i, Extended Data Fig. 3c**). Further quantitative analysis revealed a very strong correlation between age and CD8 infiltration (**Fig. 1j**). Notably, a similar pattern was observed in postmortem human hypothalamic tissues (**Fig. 1k, l**), confirming the infiltration of CD8 is conserved process between murine and human aging. CD8 in the aged hypothalamus were primarily localized near the third ventricle, median eminence, and arcuate nucleus, the regions adjacent to fenestrated capillaries (**Fig.1i**). Although, we also often observed CD8 in the more dorsal region such as dorsomedial hypothalamus (DMH) and ventromedial hypothalamus (VMH), and paraventricular nucleus (PVN) (**Extended Data Fig. 3d**).

We also detected CD8 across major anatomical compartments of the brain, including the lateral ventricular region (fimbria, subventricular zone), parenchyma (corpus callosum, dentate gyrus, hypothalamus), and leptomeningeal area (choroid plexus, subfornical organ) (**Extended Data Fig. 3e**). Regions derived from leptomeningeal tissue, including the choroid plexus and subfornical organ, showed expectedly high immune cell presence. Among parenchymal regions, except fimbria (a known immune gateway), the hypothalamus uniquely exhibited robust CD8 accumulation, indicating a selective vulnerability during aging (**Extended Data Fig. 3e**). This unique feature highlights the hypothalamus as particularly susceptible to immune cell infiltration during aging.

### Phenotypic Characteristics of CD8 Infiltrating the Aged Hypothalamus

To determine the phenotype of infiltrating CD8 in the aged hypothalamus, we projected our single-cell CD8 dataset onto a reference atlas of compound mouse tumor-infiltrating lymphocytes (TILs) using the projectTILs computational framework^24^. This analysis revealed a striking difference between young and old hypothalamic CD8 populations: while young CD8 aligned predominantly with the “Early Activation” subset, aged CD8 shifted toward an “Effector Memory” phenotype (**Fig. 2a-b**). Expression-level comparisons confirmed the fidelity of this mapping, as key transcriptional markers and effector molecules, including Tbx21 (*Tbx21*), granzyme (*Gzmk*), fas-ligand (*Fasl*), interferon-gamma (*Ifng*) showed consistent expression patterns with the corresponding reference subsets (**Fig. 2c-d).** Immunostaining confirmed that in the old hypothalamus, a substantial fraction (∼40%) of CD8 expressed T-bet (**Fig. 2e**). These findings suggest that CD8 in the aged hypothalamus acquire features of chronic activation.

**Fig. 2.**
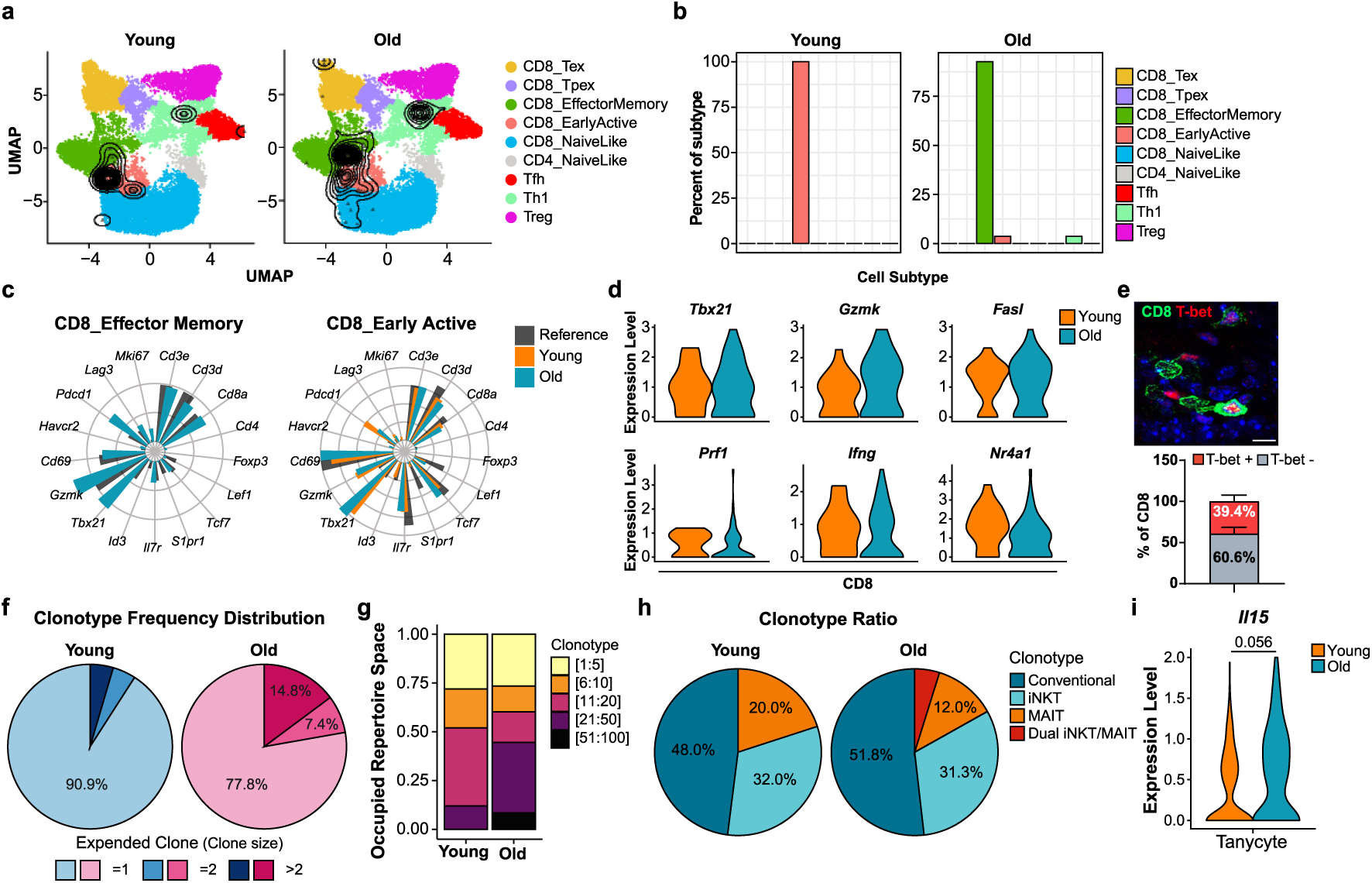
Phenotypic Characteristics of CD8 Lymphocytes Infiltrating the Aged Hypothalamus. **a-c**, Identification of T cell states by projecting CD8 in our dataset onto a reference atlas of tumor-infiltrating CD8 lymphocytes using the ProjecTILs framework. **a**, UMAP projection showing T cells from young and old mice (contour lines) overlaid on reference-defined T cell subtypes (colored clusters). Subtypes include: CD8_Tex (CD8 exhausted T cells), CD8_Tpex (CD8 progenitor exhausted-like T cells), CD8_EffectorMemory (CD8 effector/memory T cells), CD8_EarlyActive (CD8 early activated T cells), CD8_Naivelike (CD8 naïve-like T cells), CD4_Naivelike (CD4 naïve-like T cells), Tfh (CD4 follicular helper T cells), Th1 (CD4 helper type 1 T cells), and Treg (regulatory T cells). **b**, Bar plot depicting the predicted distribution of CD8 across reference-defined states. **c**, Radar plot comparing gene expression of reference states of marker genes between query cells (young: orange; old: dark cyan) and reference cells (dark gray). **d**, Violin plots showing expression of effector function genes (*Tbx21, Gzmk, Fasl, Prf1*, *Ifng*) and the antigen-stimulation marker *Nr4a1* in CD8 clusters from young and old mouse hypothalamic. **e**, Representative immunofluorescence image of CD8 and T-bet in the hypothalamus of old mice with quantification of the proportion of T-bet⁺ cells among CD8 (n = 3). Plot show mean ± s.e.m. Scale bar, 10 μm. **f-h**, TCR sequencing analysis of CD8 from young and old hypothalami. **f**, Clone size frequency distribution showing the proportion of clones across different clone size categories. **g**, Relative repertoire space of each clone group among total clones; categories are defined by sequential clone size thresholds. **h**, Proportion of clonotypes classified by TCR gene usage (Conventional, iNKT, MAIT, Dual iNKT/ MAIT). **i**, *IL-15* expression in tanycyte populations derived from scRNA-seq of young and old hypothalami. scRNA-seq derived data (d,i) statistical significance was derived from the Seurat result that is multiple testing corrected using the Benjamini and Hochberg method.

T cell receptor (TCR) sequencing showed aged hypothalamus contained a markedly increased number of detectable TCR clonotypes, with greater clonal expansion per clone compared to young controls (**Fig. 2f**), in agreement with our observation of increased CD8 infiltration (**Fig. 1i**). However, the expansion was broadly distributed across many clonotypes, with no evidence of a single dominant clone (**Fig. 2g**). Thus, while the abundance of clonally expanded CD8 increased with aging, the lack of clonal dominance argues against persistent antigen-driven expansion. Supportively, expression of NR4A1 (Nur77), a sensitive marker of persistent antigen stimulation, was unchanged in aged CD8 (**Fig. 2d**).

Further analysis revealed that nearly half of the detected clonotypes in both young and old hypothalami corresponded to unconventional T cells, as defined by atypical TCR gene usage (**Fig. 2h**). Most of highly expanded clones (frequency >5) also fell within this category. Unconventional T cells are known to respond to non-peptide antigens or be activated in innate-like fashion by cytokines such as IL-2, IL-7, IL-12, IL-15, IL-18, IFN-β, or IFN-γ^25,26^. Classical memory CD8 that infiltrate aged tissues can also undergo expansion and maintenance through cytokine-mediated bystander activation within pro-inflammatory milieus^27,28^. Analysis of our scRNA-seq dataset revealed a trend toward increased *IL-15* expression in aged tanycytes (p = 0.056) (**Fig. 2i**). Together, these results indicate that CD8 accumulation in the aged hypothalamus is unlikely to result from classical antigen-specific activation. Instead, these cells likely acquire effector memory–like features through chronic, cytokine-driven stimulation within the aged hypothalamic niche.

### Tanycytes and Microglia as Key Immune-Interacting Cell Types with Infiltrating CD8 in the Aged Hypothalamus

In our scRNA-seq data, tanycytes and microglia were identified as the cell types with the highest number of DEGs (**Fig. 3a**), suggesting they undergo the most extensive transcriptional changes during aging. This finding aligns with a previous study reporting that cell types localized near the third ventricle of the hypothalamus, including tanycytes and ependymal cells, are particularly susceptible to age-related changes^17^. Given the close developmental and spatial relationship between tanycytes and ependymal cells, it is plausible to generalize their immune response as a common tanycyte-lineage phenomenon. In parallel with CD8, GO analysis revealed that microglia, tanycytes, and ependymal cells commonly exhibited enrichment in immune-related pathways, such as ’response to interferon’, and ’peptide antigen assembly with MHC class I protein complex’ (**Fig. 3b, Extended Data Fig. 4a**).

**Fig. 3.**
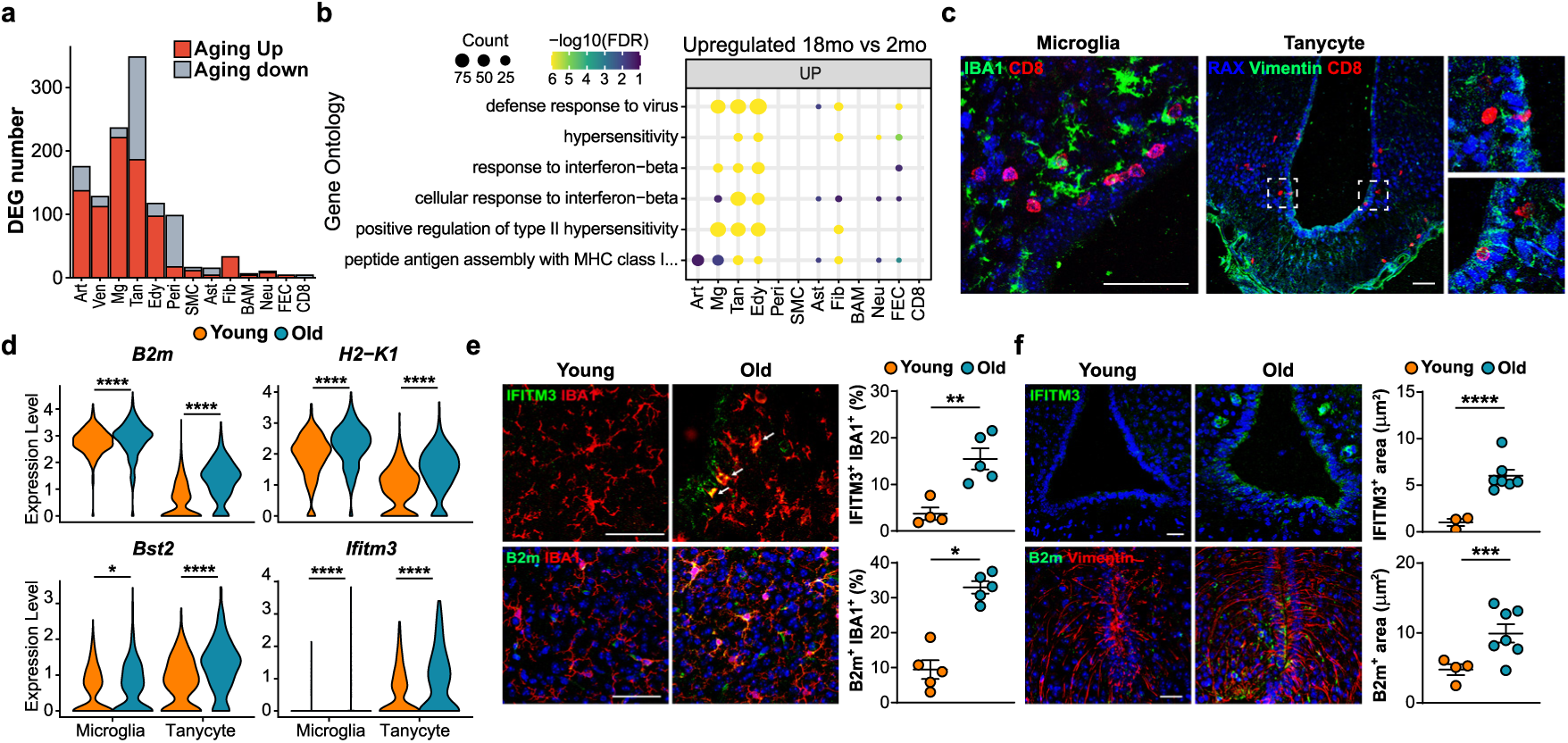
CD8 interactions with microglia and tanycytes in the aged hypothalamus. **a**, Number of DEGs per cell type (red, upregulated; gray, downregulated in old hypothalamus) from scRNA-seq data. **b**, GO analysis of DEG upregulated in the old hypothalami for each cell cluster. **c**, Representative immunofluorescence images showing CD8 in contact with IBA1⁺ microglia (left) and RAX⁺ Vimentin⁺ tanycytes (right) in aged hypothalamus. The far right panels are insets of high magnification images of CD8 and tanycyte. Scale bars, 50 μm. **d**, Violin plots showing expression of MHC class I genes (*B2m, H2-K1*) and interferon-stimulated genes (*Bst2, Ifitm3*) in microglia and tanycyte clusters from young and old hypothalamic scRNA-seq data **e-f**, Co-expression and quantification signals of IFITM3 and B2M in (e) microglia and (f) tanycytes. Quantification shows the percentage of immunoreactive cells among IBA1⁺ microglia and tanycytes identified by their localization along the third ventricle lining or Vimentin expression (n = 4∼5 mice per group). Scale bars, 25 μm. Dot plots show mean ± s.e.m. Statistical significance was determined by two-tailed Student’s t-test *P < 0.05, **P < 0.01, ***P < 0.001, ****P < 0. 0001. scRNA-seq derived data (d) statistical significance was derived from the Seurat result that is multiple testing corrected using the Benjamini and Hochberg method.

Immunohistochemical analysis revealed that microglia and tanycytes directly interact with infiltrating CD8, particularly around the third ventricle, a region with fenestrated capillaries where the absence of blood-brain barrier may facilitate immune cell entry (**Fig. 3c**). CD8 primarily target MHC-I class expressing cells and exert immune effects through interferon signaling^29^. MHC-I and interferon-stimulated genes (ISGs) expressions were significantly upregulated in microglia and tanycyte populations during aging (**Fig. 3d**). These transcriptomic findings were further validated by immunostaining for representative MHC-I (B2m) and ISG (IFITM3) proteins (**Fig. 3e-f**), confirming their close interactions with CD8. Together, these findings strongly suggest that microglia and tanycytes directly interact with infiltrating CD8, potentially driving hypothalamic aging.

### Tanycyte apoptosis by CD8 infiltrated into the aged hypothalamus

A key question is whether tanycytes or microglia act as initiators of T-cell infiltration through chemokine secretion, function mainly as responders to T-cell activity, or participate in reciprocal interactions involving both roles. Clarifying these causal relationships is essential for understanding how immune–neural crosstalk drives hypothalamic aging.

To address this, we first investigate aging-related tanycyte alterations and their interaction with CD8 by performing a detailed sub-clustering analysis of tanycytes from our scRNA-seq dataset. Our sub-clustering identified five distinct tanycyte subpopulations (**Fig. 4a**). Four clusters corresponded to established classical tanycyte subtypes, β2, β1, α2, and α1, which we validated based on known marker gene expression profiles (e.g., *Gpr50, Adm, Ptn*), which were confirmed spatially using the Allen Brain Atlas in situ hybridization dataset (**Extended Data Fig. 5a-h**). The fifth identified subcluster corresponded to tanycyte-derived neurons (TDNs), consistent with previous reports from the literature^30^. Our analysis revealed significant age-related shifts in the composition of tanycyte subtypes, characterized by a pronounced decrease in α1 tanycytes and an increase in β2 tanycytes (**Fig. 4b**). DEG analysis within these subclusters showed that the TDN subcluster exhibited the highest number of DEGs (**Fig. 4c**).

**Fig. 4.**
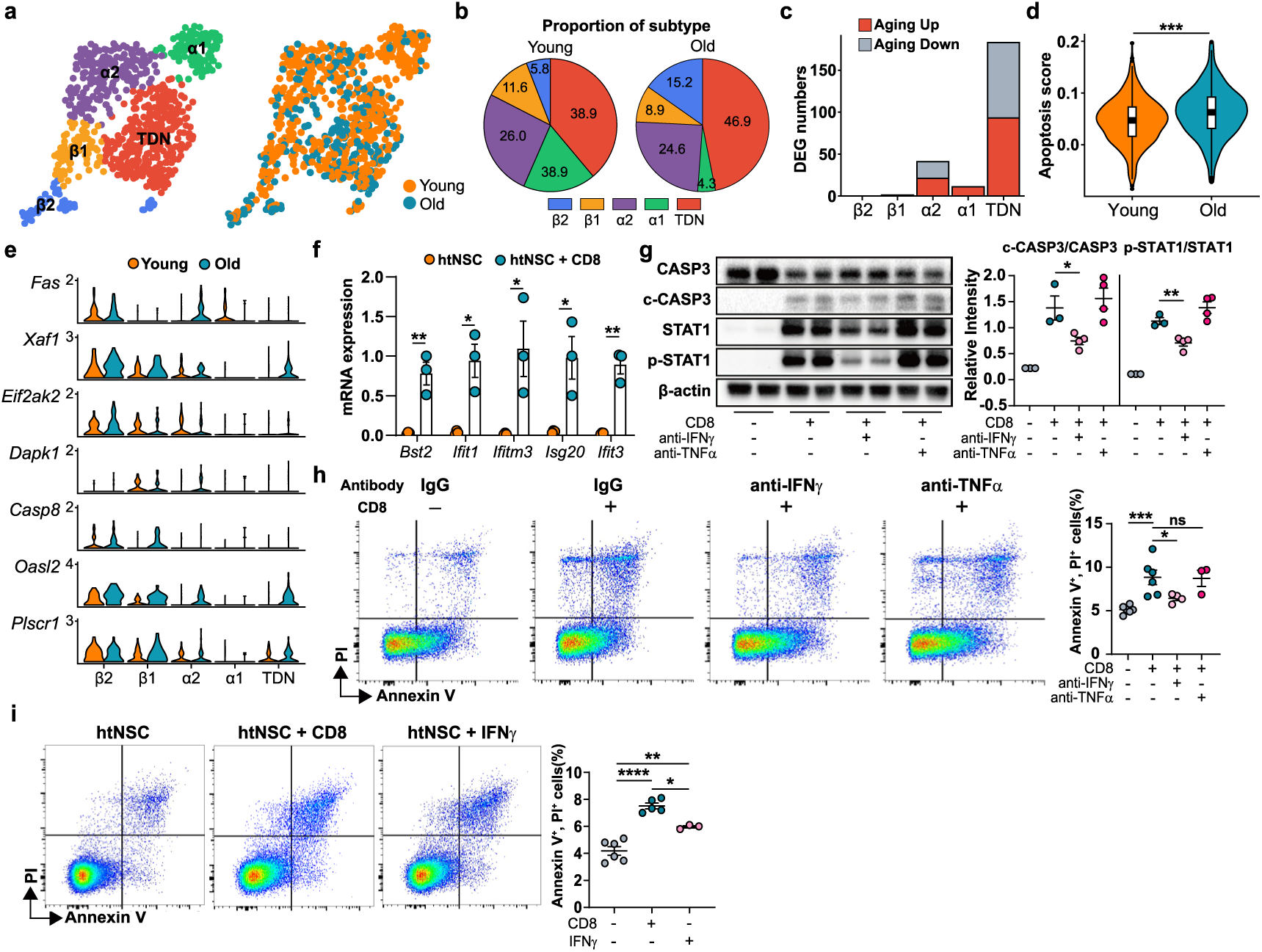
Infiltrating CD8 induces tanycytic apoptosis in the aged hypothalamus. **a–c**, Subclustering analysis of tanycytes from scRNA-seq of young and old hypothalami. **a**, UMAP projection identifying five tanycyte subtypes: β2, β1, α2, α1, and TDN (left) and the distribution of each cell by sample groups showing no batch effect. **b**, Pie charts showing the relative proportions of tanycyte subtypes in young and aged hypothalamus. **c**, Numbers of DEGs per subtype (red, upregulated; gray, downregulated in aged hypothalamus). **d-e**, Apoptosis-associated gene expression in tanycytes from scRNA-seq data. **d**, Module score based on the Reactome term Apoptosis. Box-and-whisker plots indicate the median (center line), interquartile range (box), and 1.5× IQR (whiskers). **e**, Expression of apoptosis-inducing ISGs in tanycyte subclusters showing general increasing trends. **f-i.** *In vitro* assays of CD8–mediated interferon response and apoptosis in htNSC. **f**, qRT-PCR analysis of ISG expression in htNSCs cultured alone or co-cultured with activated CD8 (n = 4 per group). **g**, Immunoblot analysis of cleaved caspase-3 and STAT1 activation (n = 3∼4 per group). **h-i**, Flow cytometry quantification of Annexin V⁺/PI⁺ assay (n = 3∼6 per group). Dot plots show mean ± s.e.m. Statistical analyses: multiple testing using the Benjamini and Hochberg method FDR < 0.001*** (d), two-tailed Student’s t-test (f) and one-way ANOVA with Tukey’s post hoc test (g–i). *P < 0.05, **P < 0.01, ***P < 0.001, ****P < 0.0001.

Previous studies have reported a general decline in tanycyte numbers during aging^6,31^, consistent with our transcriptomic data showing a ∼40% reduction in tanycyte abundance between young and old hypothalami (11.72% young vs. 6.94% old, **Fig. 1h**). GO enrichment analysis of DEGs in tanycyte subclusters revealed significant over-representation of immune response and cytotoxicity pathways (**Extended Data Fig. 5i**), suggesting that infiltrating CD8 contribute to tanycytic death an hypothalamic dysfunction. Supportively, a module score based on Reactome pathway of apoptosis was significantly elevated within aged tanycytes (**Fig. 4d**). Furthermore, ISGs that are previously identified capable of inducing apoptosis^32^, were notably upregulated in our aging tanycyte dataset (**Fig. 4e**), reinforcing the potential apoptotic mechanism triggered by infiltrating CD8.

To experimentally test this hypothesis, we used hypothalamic neural stem cells (htNSCs) as a tanycyte lineage model (**Supplementary Fig. 3a-b**). Co-culturing these htNSC with activated CD8 resulted in significantly increased cell apoptosis, as assessed by cleaved caspase-3 on immunoblot analyses (**Fig. 4g**) as well as Annexin V/PI assay (**Fig. 4h**). Since IFN-γ and TNF-α are key cytokines involved in cytotoxic T lymphocytes-mediated apoptosis, we assessed their roles by introducing neutralizing antibodies into the htNSC–CD8 co-culture. Anti-TNF-α antibody treatment had no inhibitory effect on cell death (**Fig. 4g, h**). In contrast, neutralization of IFN-γ markedly reduced CD8–induced apoptosis in htNSCs. Consistently, STAT1, key target, of IFN-γ, was activated in htNSCs following co-culture with CD8, and this activation was suppressed by anti–IFN-γ treatment (**Fig. 4g, h**). The apoptotic response was accompanied by upregulation of ISGs in htNSCs (**Fig. 4f**), consistent with our previous in vivo and scRNA-seq findings (**Fig. 3b, d**), reinforcing the involvement of IFN-γ signaling in this process. Notably, although recombinant IFN-γ alone could induce apoptosis, its effect was less potent than that of direct T cell co-culture (**Fig. 4i**), suggesting that additional T cell-derived factors may synergize with IFN-γ to enhance cell death in tanycytes. Together, these findings provide converging evidence that infiltrating CD8 directly induce apoptosis of tanycytes in the aged hypothalamus, through mechanisms partially mediated by IFN-γ signaling.

### Hypothalamic Ccl3 and Ccl4 Expression Recruits CD8 and Accelerates Aging Phenotypes

Having established that CD8 infiltrate the aged hypothalamus and induce tanycyte apoptosis, our next objective was to determine how CD8 accumulate in the aging hypothalamus. Because CXCL- and CCL-family chemokines serve as canonical chemoattractant for CD8, we examined both our bulk RNA-seq and scRNA-seq datasets for chemokine changes. Multiple transcripts were elevated in aged samples, but *Ccl3* emerged as the most consistent and significantly up-regulated chemokine. *Ccl3* was robustly increased in bulk RNA-seq and enriched in the microglia cluster of the scRNA-seq data (**Fig. 5a)**. The closely related *Ccl4* transcript followed the same pattern, albeit without reaching statistical significance **(Fig. 5a).** Notably, the CD8 cluster itself expressed high levels of *Ccr5* **(Fig. 5a**), the shared receptor for both CCL3 and CCL4, suggesting a chemokine-receptor axis capable of guiding T cell migration into the tissue. To corroborate the transcriptomic results, we quantified CCL3 protein with ELISA and observed a significant increase in aged hypothalamic lysates (**Fig. 5b**). Immunofluorescence again confirmed a ∼5-fold rise in CCL3-positive microglia in old mice (**Fig. 5c**).

**Fig. 5.**
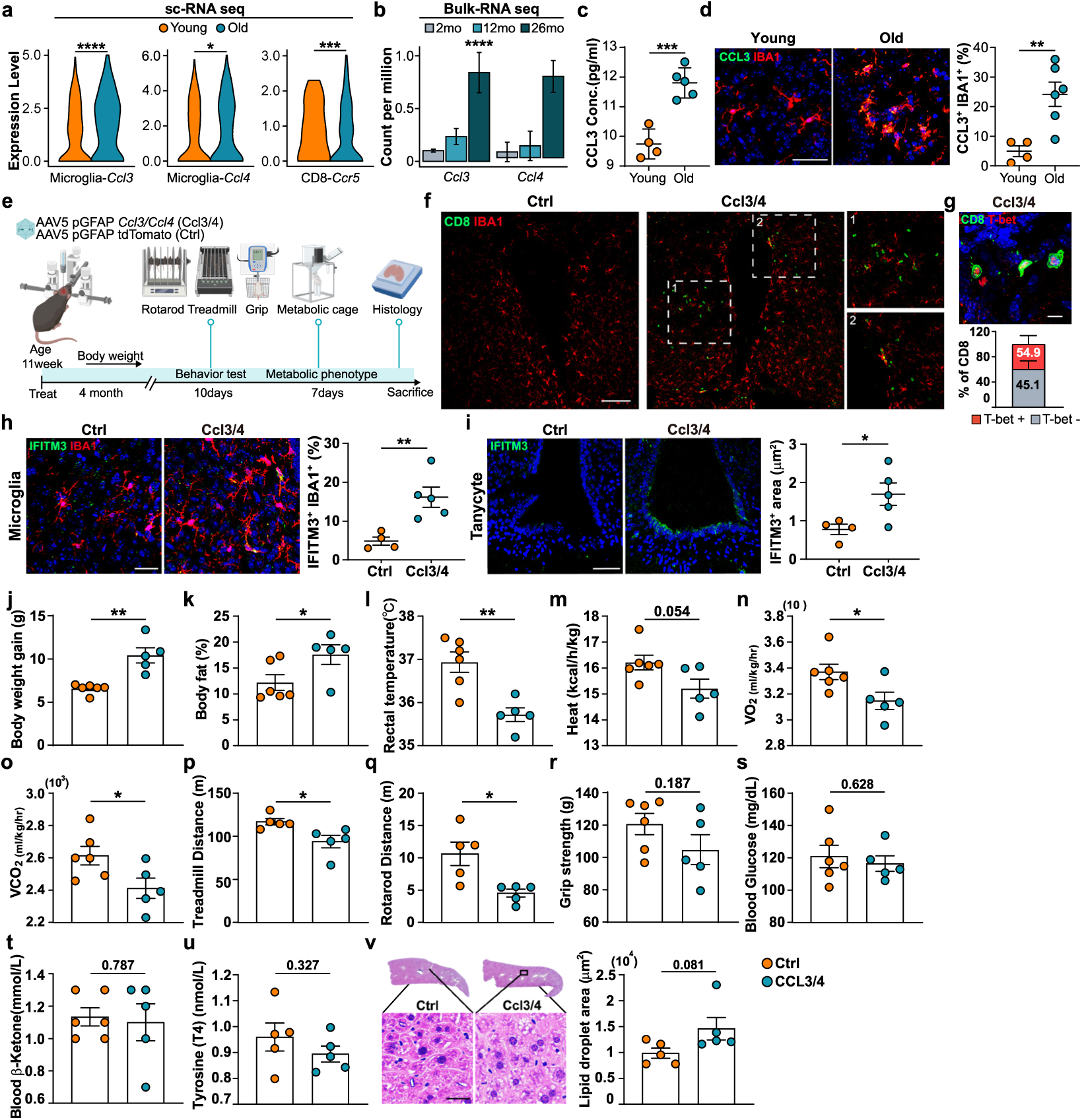
Microglia-derived CCL3/4 recruits CD8 to the aging hypothalamus and drive systemic aging phenotypes. **a**, Age-dependent increase of *Ccl3* and *Ccl4* transcripts in hypothalamic microglia shown by scRNA-seq (left, violin plots; young vs old) (left) and expression of *Ccr5* in CD8 (right) **b.** Bulk RNA-seq data shows age-dependent upregulation of *Ccl3* and *Ccl4* at 2, 12, and 26 months. **c**, ELISA detection for CCL3 protein level from young and aged hypothalamus tissue (young, n = 4; old, n =5). **d**, Representative immunofluorescence images of CCL3⁺ IBA1⁺ microglia in the hypothalamus of young and aged mice, with corresponding quantification (young, n = 4; old, n=5). Scale bars, 50 μm. **e-v**, Effects of intra-hypothalamic AAV5-pGFAP-*Ccl3/4* administration in young mice (3–months; Ctrl, n = 6; Ccl3/4, n =5). **e**, Experimental schematic. **f–i**, Immunohistochemical analysis showing infiltrated CD8 interacting with IBA1⁺ microglia and (**f**) proportion of T-bet^+^ CD8 (n = 4) (**g**), IFITM3 expression in microglia (**h**), and in tanycytes (**i**) of AAV5-pGFAP-*Ccl3/Ccl4*-injected versus control (AAV5-pGFAP-tdTomato injected) hypothalami (Ctrl, n =4; Ccl3/4, n =5). Scale bars: e, 100 μm; f, 10 μm; g, 25 μm; h, 50 μm. **j-v**, Assessment of systemic aging phenotypes 4 months after AAV5-Ccl3/4 (or AAV-Ctrl) injection, including body weight (**j**), body composition analysis (**k**), rectal temperature (**l**), energy expenditure (**m**) and O₂ (**n**) and CO₂ (**o**) consumption in metabolic cages, treadmill endurance (**p**), rotarod performance (**q**), grip strength (**r**), fasting blood glucose (**s**), blood ketone levels as represented by BHB (**t**), serum thyroxine (T4) levels (**u**), and liver histology with quantification of lipid droplet area (**v**) (Ctrl, n = 5∼6; Ccl3/4, n =5). Scale bar, 30 μm. Bars and Dot plots show mean ± s.e.m. Statistical significance was determined by two-tailed Student’s t-test (c-v), Wilcoxon (a), or DESeq2 (b). *P < 0.05, **P < 0.01, ***P < 0.001.

Building on our in vivo findings, we sought to determine whether the CCL3/CCL4–CD8 axis observed in aging hypothalami is sufficient to drive CD8 infiltration and downstream aging-like phenotype in vivo. To test this, we delivered AAV5 vectors encoding mouse *Ccl3* and *Ccl4* under the control of the astrocyte-specific GFAP promoter (AAV5-pGFAP-*Ccl3/Ccl4*) into the hypothalami of young and middle-aged mice (**Fig. 5d and Extended Data Fig. 6a**). We targeted astrocytes rather than microglia because microglia are sparse in the healthy young hypothalamus and current AAV tools for microglia-specific delivery remain limited. Although CCL3 and CCL4 in our findings are produced by microglia, astrocytic expression was expected to recreate a chemoattractant environment sufficient to drive T cell recruitment.

One month after AAV administration in young-aged mice, we observed robust recruitment of CD8 following increased expression of CCL3 in the hypothalamus (**Extended Data Fig. 6a-d**). Importantly, this was not seen in control virus-transduced animals, ruling out the possibility that the CD8 infiltration resulted from viral infection or transduction artifact (**Extended Data Fig. 6c-d**). Similar to aged mice, the CD8 were predominantly localized around the third ventricle (**Extended Data Fig. 6d**) and remained in the hypothalamus for at least five months post-injection, the longest time window assessed (**Fig. 5e**). Notably, approximately 50% of CD8 stained positive for T-bet, a transcription factor for effector T cell activation (**Fig. 5f**). This supports the idea that CD8 in the aged or *Ccl3/4-*induced hypothalamus become activated in situ without ectopic antigen presentation. Mirroring the aged hypothalamus, AAV*-Ccl3/4*-injected mice showed increased IFITM3 immunoreactivity in both microglia and tanycytes (**Fig. 5g-h**), indicating a local interferon response consistent with CD8 engagement during natural aging.

We next assessed the systemic consequences of *Ccl3/4*-driven CD8 infiltration into the hypothalamus on age-associated physiological decline. Systemic aging is marked by impaired energy and lipid metabolism together with reductions in physical capacity. In natural aging, 16–20-month-old mice exhibited declines in metabolic function, motor activity, coordination, muscle strength, and endurance compared with 3–4-month-old controls **(Extended Data Fig. 7a–k,o–p)**. In contrast, glucose tolerance and cognitive performance were largely preserved **(Extended Data Fig. 7l–n)**, consistent with reports that memory and cognition remain relatively intact until ∼22 months of age^33^. The age-dependent changes in body metabolism and behaviors were phenocopied in the young mice by the AAV*-Ccl3/4* injection. Specifically, by four months, mice in the AAV*-Ccl3/4* group exhibited significantly greater weight gain relative to controls (**Fig. 5i**), along with increased fat mass as confirmed by body composition analysis (**Fig. 5j**). This was accompanied by reduced systemic energy expenditure, including heat generation and lower oxygen consumption, carbon dioxide production (**Fig. 5k-n**). Histological examination also revealed early evidence of hepatic steatosis, although the lipid droplet accumulation did not reach statistical significance (**Fig. 5u**). Functionally, these mice displayed impaired physical motor performance, including reduced motor coordination and endurance on both the treadmill tests and rotarod (**Fig. 5o-p**). Grip strength also trended downward (**Fig. 5q**). Along with no changes of fasting blood glucose and ketone levels (**Fig. 5r-s**), plasma thyroxine (T4) levels remained unchanged, indicating that the hypothalamic–pituitary–thyroid (HPT) axis remained intact despite CD8 infiltration (**Fig. 5t**). These findings suggest that CD8–mediated hypothalamic dysfunction is likely mediated by autonomic nervous system signaling than neuroendocrine disruption.

The middle-aged cohort exhibited similar albeit less pronounced effects (**Supplementary Fig. 4a-b**). This “ceiling effect” implies that some hypothalamic dysfunction may already be present by midlife, limiting responsiveness to additional CCL3/4-induced perturbation. Next, to test whether CD8 can persist in the young hypothalamus, we transplanted them into this region. The grafted cells showed poor survival, indicating that additional chemokines, cytokines, or local cues are required for their maintenance (**Supplementary Fig. 5**).

Taken together, these results show that hypothalamic overexpression of CCL3 and CCL4 is sufficient to recruit CD8, activate local glial interferon responses, and trigger multiple systemic features of aging in otherwise healthy young mice. This provides strong in vivo evidence supporting the causal role of hypothalamic CD8 infiltration in driving both local cellular remodeling and whole-body aging phenotypes.

### Lipid-Accumulating Microglia as the Primary CCL3/4 Source in the Aged Hypothalamus

Sub-clustering of the microglia subset from our scRNA-seq dataset revealed three distinct microglial populations (**Fig. 6a**) including ‘homeostatic microglia’, expressed canonical markers such as *P2ry12*, *Cx3cr1*, and *Tmem119*, consistent with previous studies^34^ (**Fig. 6b**). The second cluster, which we term ‘activated microglia’, showed high expression of integrated stress response (ISR) genes (*Atf3*, *Atf4*, *Ddit3*). Given uncertainties about whether this population reflects a bona fide biological state or an artifact from single-cell isolation, we adopted the broad “activated” terminology from Marsh, et al.^35^. The third population corresponded to ‘lipid droplet–accumulating microglia (LDAM)’, characterized by *Plin2*, *Plin3*, and *Apoe* expression as described previously^36^ (**Fig. 6b**).

**Fig. 6.**
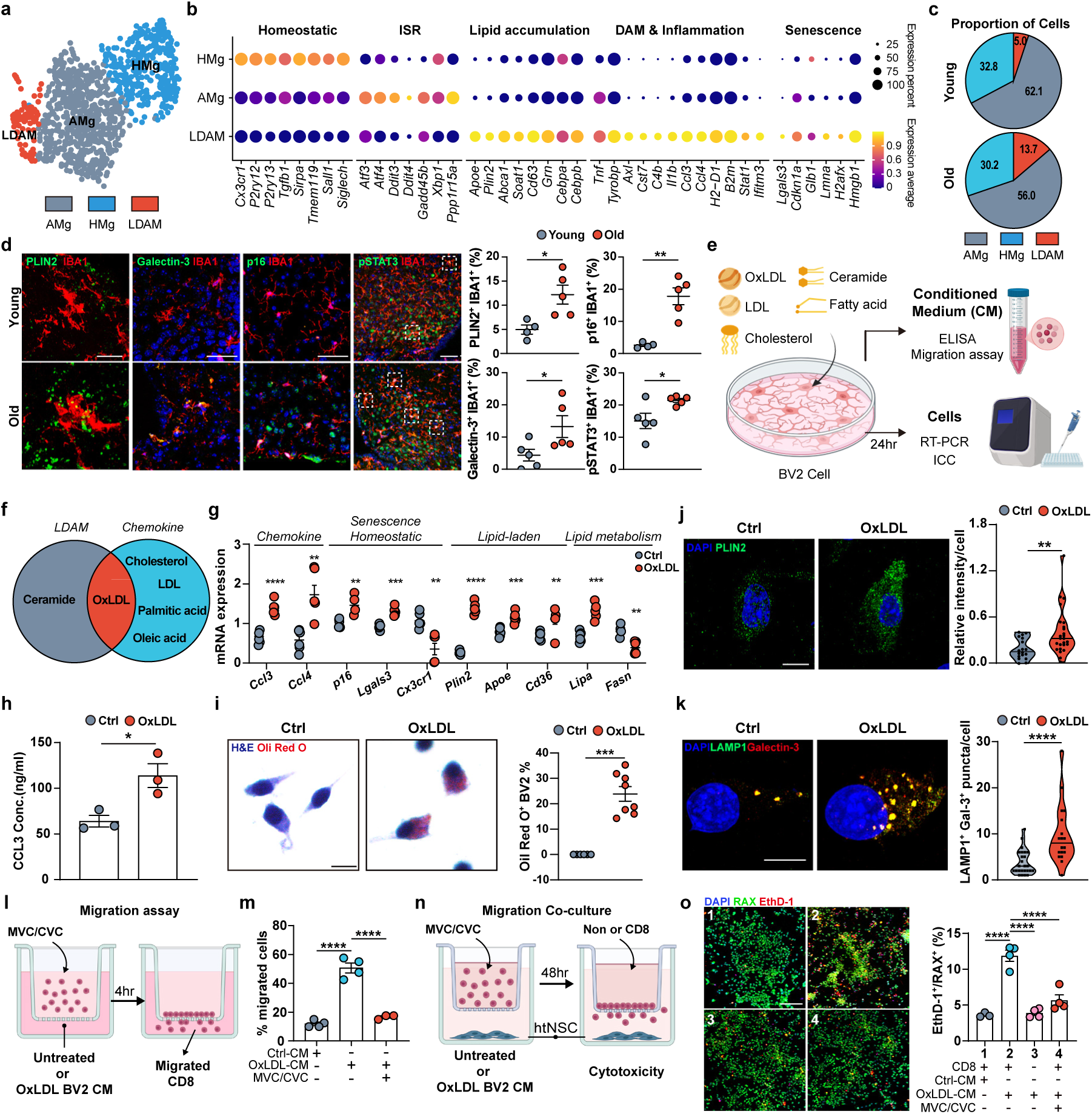
Oxidized LDL drives lipid droplet–accumulating microglia (LDAM), which recruit CD8 to the aging hypothalamus via the Ccl3/4–CCR axis. **a–c**, Subclustering analysis of microglia from young and aged mouse hypothalami. **a**, UMAP projection of identified three transcriptionally distinct subtypes: activated microglia (AMg), homeostatic microglia (HMg) and lipid-accumulating microglia (LDAM). **b**, Dot plots for cluster marker gene expressions of microglia subcluster. **c**, Pie charts indicating the relative proportion of each subtype across age groups. **d**, Immunohistochemistry of hypothalamic IBA1⁺ microglia expressing LDAM markers PLIN2, Galectin-3, p16, and pSTAT3 in young and aged mice (n = 4∼5 per group). Scale bars: 10 μm (PLIN2), 50 μm (Galectin-3, p16, pSTAT3). **e-j**, Induction of the LDAM phenotype by OxLDL in BV2 microglia. **e**, Schematic of the lipid screening assay for LDAM induction and CCL3 secretion. **f**, Venn diagram summarizing the effect of screened lipid species **g,** qRT–PCR analysis of genes related to chemokine signaling (*Ccl3, Ccl4*), senescence (*p16, Lgals3*), homeostasis (*Cx3cr1*), lipid accumulation (*Plin2, Apoe, Dgat1, Cd36*), and lipid metabolism (*Pnpla2, Lipa, Fasn*) (Ctrl vs. OxLDL; n = 5∼6). **h**, ELISA measurement of secreted CCL3 protein in conditioned media (Ctrl vs. OxLDL; n = 3). **i**, Oil Red O staining showing intracellular lipid droplet accumulation (Ctrl: n = 6; OxLDL: n = 8 cells). **j**, Immunofluorescence for PLIN2 expression (Ctrl: n = 18; OxLDL: n = 25 cells). **k**, Detection of lysosomal damage marked by LAMP1⁺Galectin-3⁺ puncta (Ctrl: n = 25; OxLDL: n = 18 cells). Scale bars: 10 μm. **l-m**, Schematic (l) and quantification (m) for transwell migration assay of CD8⁺ T cells toward conditioned medium (CM) from untreated BV2 cells (Ctrl-CM), OxLDL-treated BV2 cells (OxLDL-CM) only, or OxLDL-CM and CD8 in the upper chamber ± MVC/CVC after 4 h (n = 3 per group). **n-o**, Migration Co-culture assay. **n**, Schematics showing htNSC were cultured for 48 h with Ctrl-CM or OxLDL-CM in the lower chamber and CD8 or not in the upper chamber ± MVC/CVC. **o**, EthD-1⁺RAX⁺staining and quantification for percent EthD-1⁺RAX⁺ cells (n = 3∼4 per group) (n). Scale bar, 100 µm. Dot plots show mean ± s.e.m. Statistical significance was determined using two-tailed unpaired Student’s t-test or one-way ANOVA with Tukey’s post hoc test. *P < 0.05, **P < 0.01, ***P < 0.001, ****P < 0.0001.

LDAM displayed additional features relevant to immune activation and senescence. They expressed inflammation-adjacent profiles including disease-associated microglia (DAM)-like markers (*Tyrobp*, *Axl*, *Cst7*), interferon response genes (*B2m*, *Stat1*, *Ifitm3*), and senescence-associated genes such as *Cdkn1a* (p21), *Cdkn2a* (p16), and *Lgals3* (galectin-3). Notably, LDAM showed the highest *Ccl3* and *Ccl4* expression among microglial subtypes (**Fig. 6b**), implicating them as the likely main source of the chemokines driving CD8 infiltration into the old hypothalamus. Quantitatively, LDAM proportion increased 2.7-fold in aged compared to young hypothalami (**Fig. 6c**). Immunostaining for PLIN2, a lipid droplet-associated protein, confirmed a similar increase in LDAM in aged brains (**Fig. 6d**). Immunostaining of senescence markers, including p16 and Galectin-3, were readily detected in aged hypothalamic microglia but almost absent in young microglia (**Fig. 6d**). Following senescence reporting guidelines^37^, we also examined pSTAT3 as a marker of pro-secretory phenotype: aged microglia showed an increased proportion of pSTAT3⁺ cells (**Fig. 6d**).

### Oxidized LDL Drives LDAM and Trigger CD8 Infiltration in the Aging HypothalamusWe next sought to determine how the lipid-laden microglia phenotype arises in the aged hypothalamus

One possible scenario is that infiltrating CD8 induce this LDAM phenotype, thereby establishing an autoregulatory loop that recruits additional T cells via the CCL3/4–CCR5 axis. However, in our *Ccl3/4*–overexpression mouse model, robust hypothalamic recruitment of CD8 did not induce LDAM-associated phenotypes or senescence-associated markers such as PLIN2 and Galectin-3, p16 (**Extended Data Fig. 8a**). Furthermore, co-culturing primary microglia with activated CD8 failed to induce the expression of LDAM-specific genes **(Extended Data Fig. 8b**). By contrast, pro-inflammatory genes (*Tnf*, *Ifng*) were strongly upregulated by co-culture, suggesting that while infiltrating CD8 can promote a pro-inflammatory microglial phenotype, they are not responsible for the LDAM features associated with lipid accumulation, senescence, or CCL3/4-mediated T cell recruitment in the aged hypothalamus.

In our dataset, transcript associated with de novo lipid biosynthesis of microglia and module scores for the GO term “neutral lipid biosynthetic process” (GO:0046460) in any cell type showed no significant increase (**Extended Data Fig. 9a-b**), making in situ lipid biosynthesis is an unlikely explanation. Given that hypothalamic microglia are positioned to sense circulating lipid signals and are prone to lipid-droplet accumulation^38–40^, and that plasma lipid species, including ceramide, free fatty acids, cholesterol, LDL, and modified LDL increase with age^41–43^, we asked whether lipid droplets in LDAM microglia might arise from uptake of circulating lipids. We screened multiple lipid species for their ability to induce upregulation for CCL3 expression and LDAM phenotype in BV2 microglia and found that only oxidized LDL (OxLDL) simultaneously increased CCL3 protein expression and LDAM gene markers. (**Fig. 6e-f and Extended Data Fig. 9c-d**). ELISA further confirmed elevated CCL3 release into the culture medium following OxLDL treatment (**Fig. 6h**). OxLDL–treated microglia recapitulated key transcriptional and phenotypic features of aged hypothalamic LDAM, including upregulation of *Ccl3*/*Ccl4*, downregulation of homeostatic markers, increased expression of senescence-related and lipid droplet–associated genes, enhanced lipid catabolism, and reduced lipid biosynthesis (**Fig.6g**). Phenotypically, they exhibited lipid-droplet accumulation validated by Oil Red O staining and PLIN2 immunostaining. (**Fig. 6i-j**), as well as LAMP1⁺ Galectin-3⁺ lysosomal vesicle accumulation, indicative of autolysosomal dysfunction (**Fig. 6k**).

Consistent with prior reports^44,45^, plasma from aged mice contained significantly higher levels of OxLDL than that from young controls (Extended Data Fig. 9e). OxLDL binds to microglial scavenger receptors and is taken up excessively in an unregulated manner^46^. In our scRNA-seq data, LDAM in the aged hypothalamus expressed elevated levels of lipoprotein scavenger receptors, including *Apoe*, *Scarb1*, *Cd44*, and *Lrp1* (**Extended Data Fig. 9f**). Furthermore, module scores for “Cellular response to low-density lipoprotein particle stimulus” (GO:0071402) and “Cellular response to oxidized LDL particle stimulus” (GO:0140052) were significantly increased in aged hypothalamic microglia (**Extended Data Fig. 9g**), supporting the physiological relevance of OxLDL–driven acquisition of the LDAM phenotype in the aged hypothalamus. Moreover, PLIN2 was markedly enriched in the hypothalamus relative to other brain regions (**Supplementary Fig. 6**), consistent with regional lipid-droplet accumulation and a heightened susceptibility to LDAM-induced CD8 recruitment.

To test whether OxLDL–treated microglia promote CD8 infiltration, we performed a transwell migration assay (**Fig. 6l**). Conditioned medium (CM) from OxLDL–treated BV2 cells markedly increased CD8 migration relative to CM from untreated cells (Ctrl-CM), and this effect was abolished by CCR5 antagonists maraviroc (MVC) and cenicriviroc (CVC), confirming the requirement for CCL3/4–CCR5 signaling in this process (**Fig. 6m**). We then asked whether blocking CCR5-dependent migration protects htNSCs from cytotoxicity of CD8. htNSCs were plated in the lower chamber with CM from OxLDL–treated BV2 cells (OxLDL-CM) or untreated CM (Ctrl-CM), and CD8 were added to the upper chamber (**Fig. 6n**). Along with T cell migration to the lower chamber (**Fig.6l**), OxLDL-CM promoted htNSC death, as shown by Ethidium homodimer-1 (EthD-1) staining; treatment with MVC and CVC in the lower chamber significantly reduced this cytotoxicity (**Fig.6o**). Notably, OxLDL-CM alone (no T cells) did not induce htNSC death (**Fig.6o**). Taken together, age-dependent OxLDL uptake transforms hypothalamic microglia into LDAM, which secretes CCL3/4 and drives CCR5-dependent recruitment of CD8s. Consistently, analysis of the human hypothalamic snRNA-seq dataset HYPOMAP^47^, revealed a significantly increased population of *CCL3⁺ PLIN2⁺ APOE⁺* microglia in aged individuals (>65) compared with younger controls (<65) (**Extended Data Fig 9h**).

### Pharmacological CCR5 Blockade in Aged Mice Prevents Hypothalamic CD8 Infiltration and Restores Systemic Physiology

We next asked whether pharmacological CCR5 inhibition could prevent T cell accumulation and attenuate aging-associated physiological decline. CCR5 antagonists have been extensively studied clinically, and some are in approved use with favorable safety profiles, making this pathway an attractive therapeutic target.

Although MVC is safe and effective in humans, our pilot tests showed incomplete and inconsistent inhibition in our murine model (data not shown), consistent with reports of its reduced potency against murine CCR5^48,49^. By contrast, CVC has demonstrated efficacy in neurological contexts in murine model^50^. To maximize inhibition in our murine model, we employed dual blockade with MVC and CVC. We next compared systemic versus central delivery in the AAV*-Ccl3/4* model. Intracerebroventricular infusion via osmotic pump nearly abolished CD8 infiltration, while intraperitoneal administration produced a comparable reduction **(Extended Data Fig. 10a)**, suggesting that peripheral delivery is a viable alternative. Based on these results, we administered MVC and CVC intraperitoneally for two months, starting at 16 months of age **(Fig. 7a)**.

**Fig. 7.**
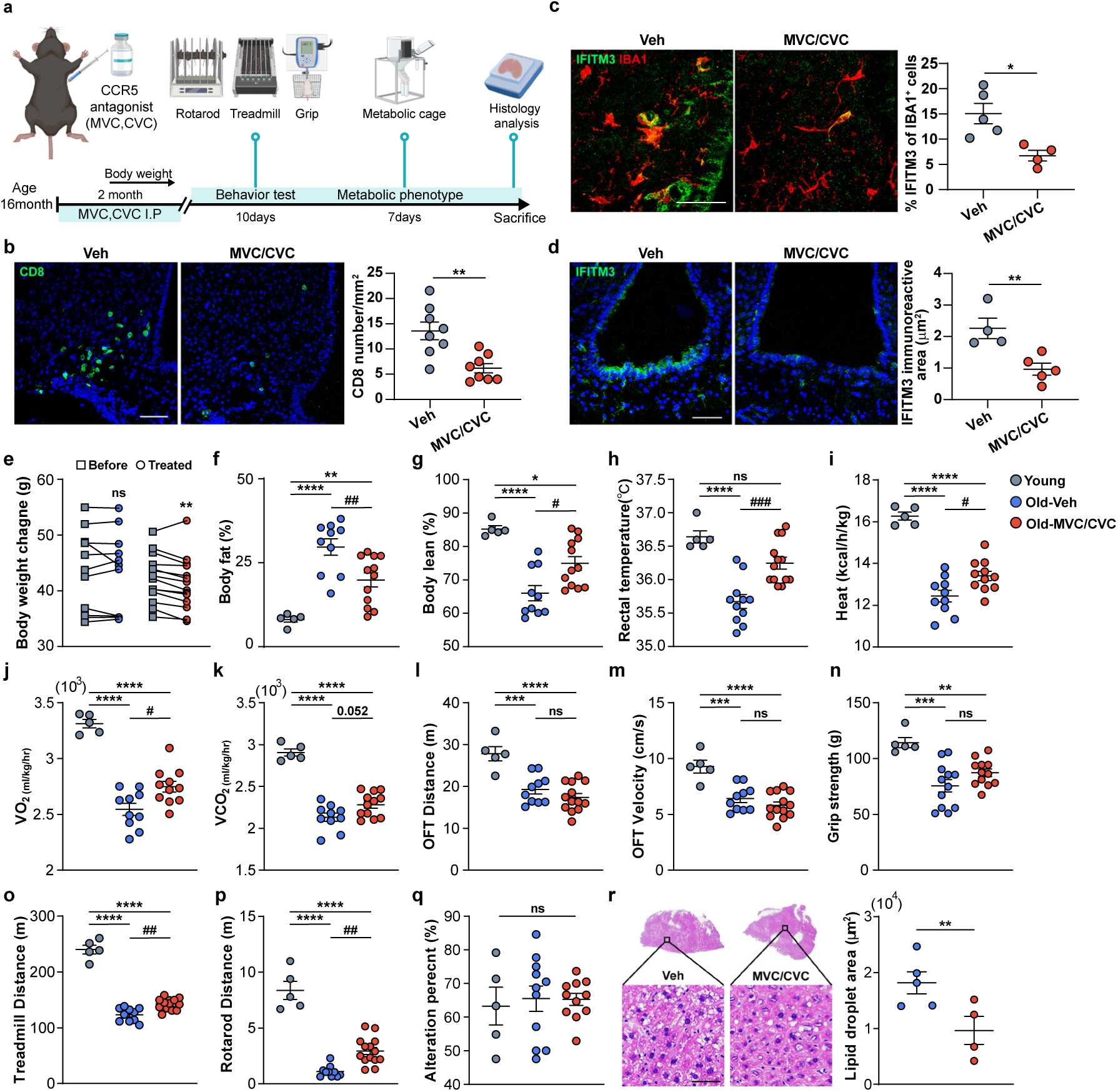
CCR5 inhibition prevents intrahypothalamic T cell infiltration and alleviates systemic aging phenotypes. **a**, Experimental workflow of CCR5 antagonist (MVC/CVC, n =14) or vehicle (n = 11, Cont) intraperitoneal (I.P) injection, histological analysis of hypothalamus and body-phenotype assessment and in mice. **b–d**, Immunohistochemical analyses of hypothalamic sections from vehicle or MVC/CVC mice. **b**, CD8 and population on hypothalamus area (n =8 Veh; n = 8 MVC/CVC). **c**, IFITM3⁺IBA1⁺ microglia and quantification for double positive microglia (n =5 Veh; n = 4 MVC/CVC). **d**, IFITM3 immunoreactivity in tanycytes lining the third ventricle (μm^2^) (n =4 Veh; n = 5 MVC/CVC). Scale bars: b-d, 50 μm. **e-q**, Body phenotype (e-h) and metabolism (i-k), and motor performance (l-p) of young and old (Veh, MVC/CVC group) (n = 5 young; n = 10∼11 Veh; n = 11∼14 MVC / CVC). Body weight gain prior and after CCR5 antagonist treatment (e), body composition analysis (f,g), rectal temperature (h), energy expenditure (i), O_2_ and CO_2_ consumption (j,k), open field test (l,m), grip strength (n), treadmill endurance (o), rotarod performance (p), Y-maze (q). **r**, Representative H&E-stained liver sections from Veh or MVC/CVC mice (n =5 Veh; n = 4 MVC/CVC) and quantification of lipid droplet area per field (μm²). Scale bars, 50 μm. Dot plots show mean ± s.e.m. Statistical significance was determined by two-tailed unpaired Student’s t-test (b-d, f) and one-way ANOVA with Tukey’s post hoc test (e-q). *P < 0.05, **P < 0.01, ***P < 0.001, ****P < 0.0001.

CCR5 inhibition–mediated reduction of CD8 infiltration into the hypothalamus was accompanied by attenuated interferon signaling, reflected by decreased IFITM3 expression in both microglia and tanycytes (**Fig. 7b-d**). This local immunomodulation translated into meaningful physiological improvements. MVC/CVC-treated mice exhibited reduced body weight gain (**Fig. 7e**), characterized by a decrease in fat mass and an increase in lean mass, accompanied by reduced hepatic lipid accumulation (**Fig. 7f-g, r**). Metabolic chamber assessments revealed increased energy expenditure, with higher rectal temperature (**Fig. 7h**), elevated heat generation (**Fig. 7i**), and enhanced oxygen consumption and CO₂ production (**Fig. 7j, k**). No improvements in locomotor activity (Open field test, **Fig. 7l, m**) and fore-limbs grip strength (Grip test, **Fig. 7n)**, and cognitive function (Y-maze, **Fig. 7q**) were observed in antagonist-treated mice. Along with the improved body energy metabolism, age-dependent decline in muscle endurance (Treadmill test, **Fig. 7o**) and motor coordination (Rotarod test, **Fig. 7p**) were significantly improved following treatment. These systemic effects support a model in which hypothalamic CD8 infiltration drives metabolic and motor decline during aging, and CCR5 blockade alleviates this trajectory.

Together, our analyses reveal that CD8 infiltrating the aged hypothalamus acquires effector memory–like phenotype, probably through non-antigen-driven, cytokine-mediated activation. Their recruitment is orchestrated by an OxLDL–LDAM–CCL3/4–CCR5 signaling axis, and pharmacological inhibition of CCR5 is sufficient to block CD8 infiltration, reduce local interferon responses, and improve age-associated decline of systemic body metabolism and motor behaviors. These findings not only establish CD8 infiltration as a proximal cause of hypothalamic aging but also highlight CCR5 inhibition as a tractable therapeutic avenue to slow neuroimmune-driven aspects of systemic aging.

## Discussion

Aging is marked by systemic metabolic dysregulation and progressive loss of homeostatic balance. Given the hypothalamus’s central role in coordinating metabolism and endocrine function, it is reasonable to propose that hypothalamic dysfunction contributes causally to aging phenotypes. Here, we identify CD8 infiltration as a prominent and defining feature of the aged hypothalamus. Importantly, this finding has a strong translational relevance since CD8 infiltration strongly correlated with chronological age in postmortem human hypothalamic tissues.

These infiltrating CD8 displayed transcriptional signatures of activation, tissue residency, and effector function, and engaged closely with tanycytes and microglia. This interaction led to tanycyte cell death and neuroinflammation. This phenomenon reflects a broader feature of immunosenescence: the gradual loss of adaptive immune competence accompanied by acquisition of heightened pro-inflammatory and cytotoxic activity. Traditionally, brain inflammation during aging and neurodegeneration has been attributed mainly to glial dysfunction and innate immune activation. However, the presence of immunosenescent T cells in aged^18,19^ and diseased brains^20,21,51^ requires a revision of this paradigm to incorporate adaptive immune cells as critical contributors to CNS aging. In addition to neuroinflammation, tanycyte loss has been observed in the aged hypothalamus and is linked to systemic physiological decline^6,52^. In this study, we demonstrate that activated CD8 drive apoptotic death of tanycytes via IFN-γ signaling, identifying T cell infiltration as the cause of tanycyte loss in aging. These findings provide a mechanistic basis for tanycyte attrition and establish CD8 infiltration as a distal driver of hypothalamic aging and systemic deterioration.

A key question is whether aging is associated with a specific “autoantigen”. Our TCR sequencing and gene expression analyses argue against this possibility. While aged hypothalami showed increased clonotype numbers and moderate expansion, no dominant clones emerged. Instead, about half of the detected clonotypes corresponded to unconventional T cells, which often recognize non-protein antigens or can be activated in antigen-independent, cytokine-driven ways^53,54^. Similarly, classical tissue-resident memory CD8 can undergo bystander activation in response to cytokines^27,28^. Together with the absence of NR4A1 induction in CD8, these findings suggest that infiltrating CD8 in the aged hypothalamus are not antigen-driven but instead activated through innate-like, cytokine-dependent mechanisms.

We further identified that microglia in the aged hypothalamus upregulate and release CCL3/4 and recruit peripheral CD8 via the CCL3/4-CCR5 axis. The CCL3/4-expressing microglia in the aged hypothalamus were characterized by lipid-droplet accumulation. Hypothalamic CD8 infiltration has been reported in obese individuals^55^, suggesting convergence between metabolic and chronological aging. Mechanistically, we implicate oxidized LDL (OxLDL) as a trigger for this process as OxLDL level rises with age^56–58^. Furthermore, lipid-laden microglia expresses scavenger receptors for OxLDL. This mirrors the cascade of atherosclerosis, where macrophages transform into pro-inflammatory foam cells via OxLDL uptake. In vitro, OxLDL treatment was sufficient to drive microglial lipid droplet accumulation, senescence, and CCL3/4 secretion, establishing a direct mechanistic link between systemic lipid metabolism and hypothalamic immune remodeling.

Ectopic expression of CCL3/4 in young mouse hypothalami was sufficient to recruit CD8 and induce systemic aging-like phenotypes. Conversely, pharmacological blockade of CCR5 with MCV/CVC abolished T cell infiltration, reduced local interferon signaling, and ameliorated systemic metabolic decline. MVC is already clinically approved for R5-tropic HIV treatment and is under evaluation in graft-versus-host disease and oncology, with a favorable safety profile. Naturally occurring CCR5 loss-of-function alleles such as CCR5Δ32 further demonstrate that CCR5 inhibition is well tolerated without serious adverse effects. We used MVC/CVC combination in this study because MVC alone only has moderate activity against murine CCR5; however, in the human context MVC alone is likely sufficient. Thus, targeting CCR5 may represent a clinically feasible healthspan promoting strategy, though benefits and risks in elderly individuals require careful evaluation.

These insights highlight clear avenues for human translation. As described, we observed age-dependent CD8 infiltration in postmortem human hypothalamic tissues. CD8-targeted PET probes, currently under development, could enable longitudinal human studies to track hypothalamic T cell infiltration and its relationship to aging biomarkers, biological aging clocks, or MRI-derived hypothalamic volume. Stratifying human cohorts by diet or circulating OxLDL levels may help resolve links between lipid metabolism, hypothalamic immune infiltration, and biological aging.

In summary, our study identifies a hypothalamus-centered mechanism of systemic aging, wherein the recruitment of CD8 and their interactions with tanycytes and microglia in the hypothalamus play a pivotal role in driving age-related physiological decline. Furthermore, we propose that therapeutic blockade of the CCL3/4–CCR5 signaling axis represents a promising strategy for mitigating immune-mediated hypothalamic dysfunction and potentially delaying systemic aging. Our study, however, has a few notable limitations. All experiments were conducted in male C57BL/6J mice, leaving sex- and strain-specific differences unaddressed. Although no sex difference in hypothalamic CD8 infiltration was found in obese humans^55^, validation in female mice remains necessary. Adaptive immunity is also heavily influenced by the microbiome, and our semi-SPF housing does not fully recapitulate human microbial diversity. These caveats highlight the importance of extending findings across models and contexts.

## Method

Extended descriptions of all experimental procedures, including in vivo mouse experiments, cell culture, molecular assays and sequencing analyses, are provided in the Supplementary Methods.

### Laboratory animal study

All animal experiments were approved by the Institutional Animal Care and Use Committee (IACUC) of the College of Medicine, Hanyang University (Seoul, Korea) under approval numbers 2021-0259A, 2023-0024A, 2023-0321A, and 2025-0002A, and were conducted in accordance with NIH guidelines. C57BL/6J male mice were used at different ages: young (7–20 weeks), middle-aged (11–15 months), and old (16–24 months). Middle-aged and old mice were purchased from the Korean Basic Science Institute (Gwangju, Korea), while young mice, pregnant females (E12), and P1 pups used for primary glial and hypothalamic neural stem cell cultures were obtained from DBL (Seoul, Korea). All mice were housed in a specific pathogen-free facility under a 12-hour light/dark cycle (08:00–20:00) with ad libitum access to irradiated standard chow (5053 PicoLab Rodent Diet 20, Labdiet, USA) and water.

### Human postmortem brain sample

Postmortem human hypothalamic brain tissue was obtained from the Dementia Brain Bank of Seoul National University Hospital and the Boston University Alzheimer’s Disease Research Center with approval from the IRB of Hanyang University (HYUIRB-202502-004). Informed consent for participation and brain donation was obtained from the donors’ legal representatives. All donors were male; three were young (22–25 years) and seven were older (60–82 years). Donor-level information is provided in Supplementary Table 1.

### Statistical analysis

Statistical analyses were performed using GraphPad Prism software (version 8; GraphPad Software) and R software. All data are presented as mean ± standard error of the mean (s.e.m.). Normality of data distributions was assessed using the Shapiro–Wilk test. For normally distributed data, two-tailed unpaired Student’s t-tests were used to compare two groups, and one-way ANOVA followed by Tukey’s multiple comparisons test was used for multiple group comparisons. For non-normally distributed data, the Mann–Whitney U test was applied. Paired t-tests were used when comparing matched measurements from the same subjects. Pearson’s correlation coefficient was used to assess linear relationships between variables. P-values less than 0.05 were considered statistically significant. All transcriptomic statistical procedures, including DEG and enrichment analyses, are described separately in the respective sections.

## Data availability

Data availability. The bulk RNA-seq and single-cell RNA-seq datasets generated in this study are available in the Gene Expression Omnibus (GEO) under accession GSEXXXXX. Source Data underlying the figures are provided with this paper. De-identified human postmortem metadata are shared as permitted by IRB/brain-bank policies. External Validation sets : human HYPOMAP snRNA-seq dataset of young and old human hypothalamus for Lipid accumulated microglia (John A Tadross et al. PMID: 39910307).

## Code availability

Analysis and figure-generation scripts for **bulk RNA-seq** and **single-cell RNA-seq** are available on **GitHub** : xxxxxxxx

---

**Extended Fig. 1.**
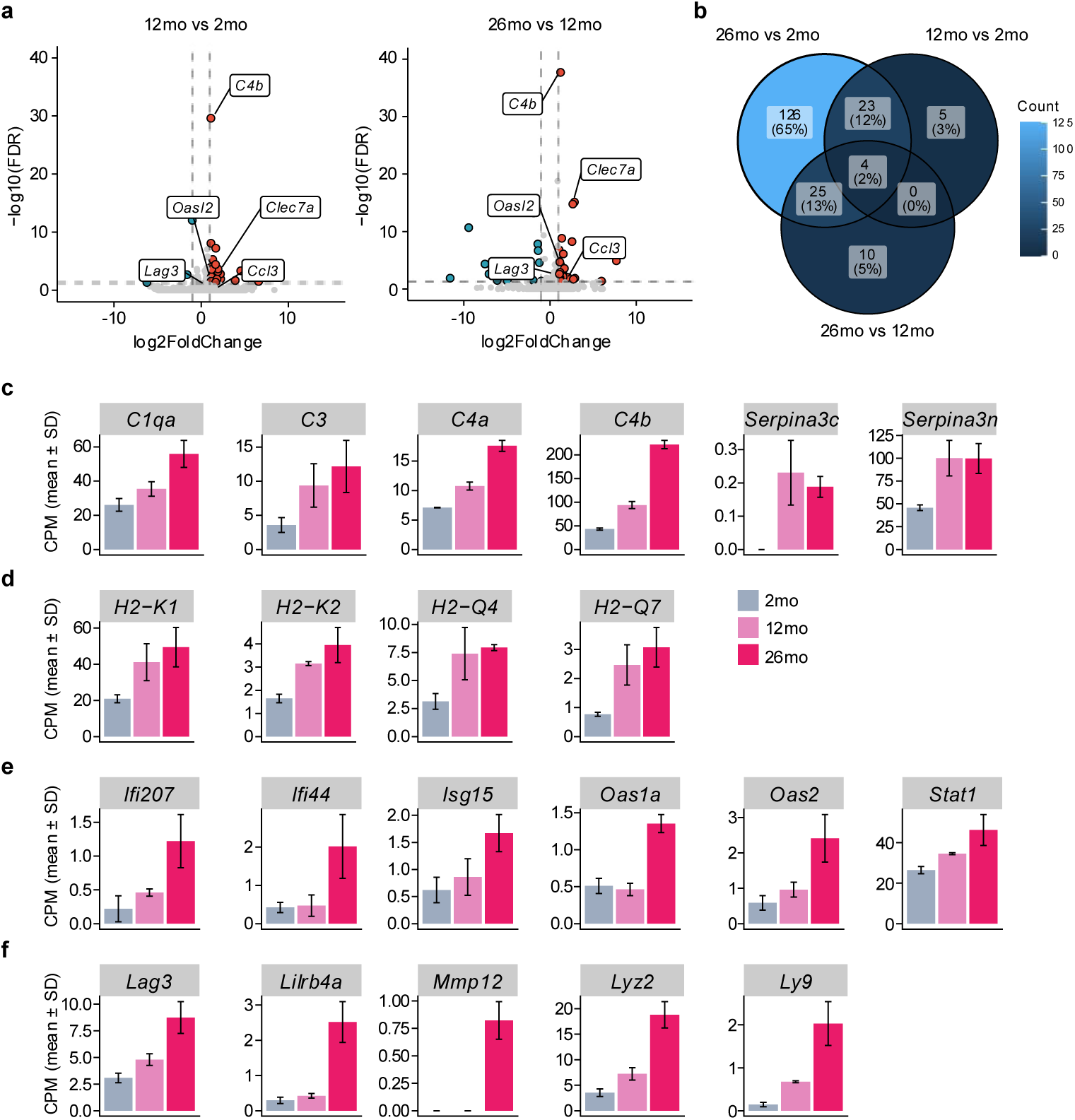
Bulk RNA-seq analysis of immune-related signatures with distinct aging patterns in the hypothalamus. **a** 2,6-Volcano plots of differentially expressed genes (DEGs) in 12-month vs 2-month mice and month vs 12-month mice. **b**, Venn diagram showing shared and unique DEGs among -three pairwise comparisons: 12-month vs 2-month, 26-month vs 2-month, and 26-month vs 12 month hypothalamic tissues. **c-f**, Bar plots of counts per million (CPM) for representative transcripts related to the complement cascade and Serpina-family protease inhibitors (c), MHC class I molecules (d), interferon-stimulated genes (ISGs) (e), and lymphocyte immune response (f).

**Extended Fig. 2.**
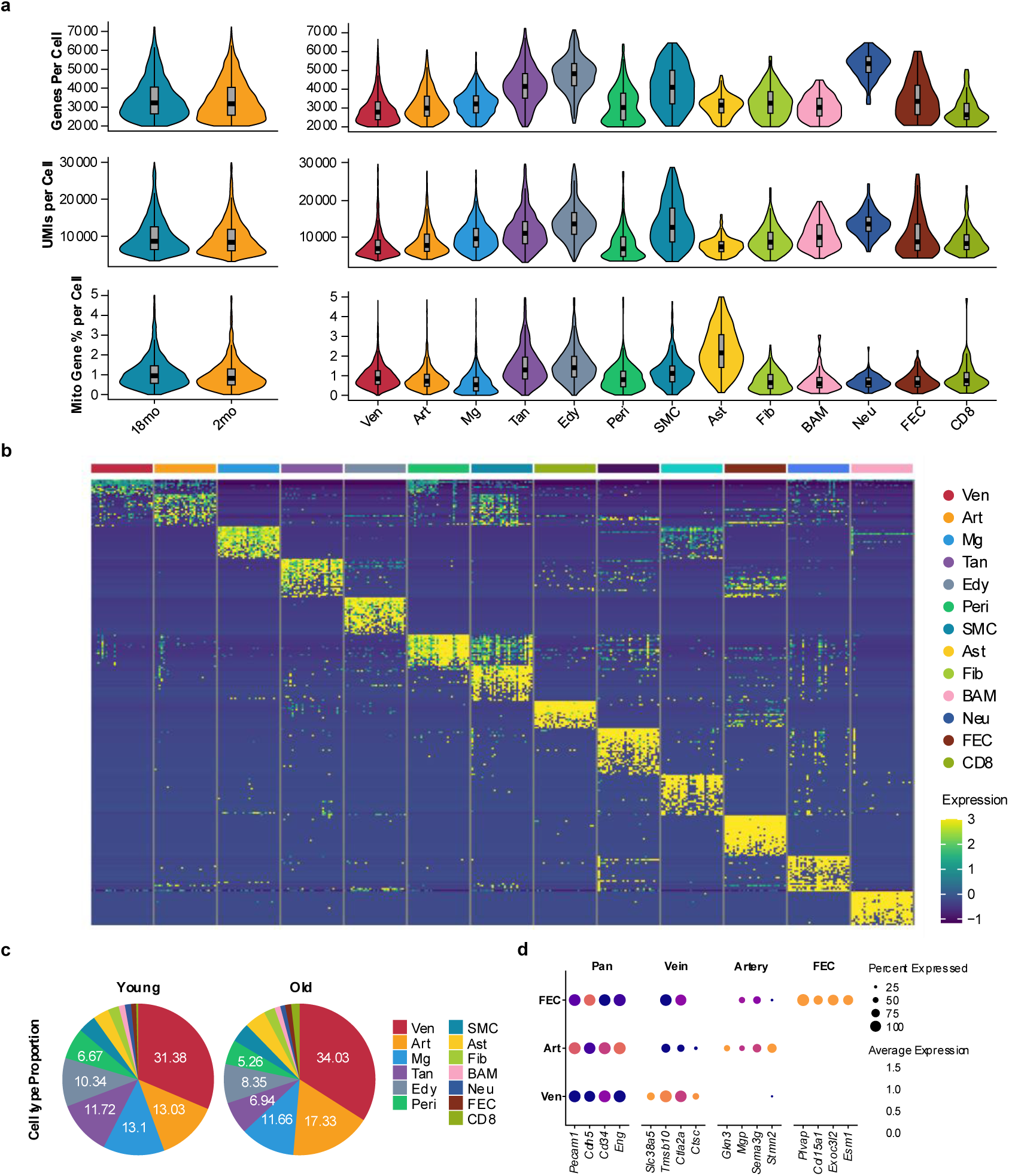
Quality control for scRNA-seq data set and transcriptomic identities of cell subtypes annotated from the scRNA-seq data. **a**, Quality control of the sc-RNA-seq dataset showing gene count, UMI count, and mitochondrial gene percentage per cell across age groups (left) and cell-type clusters (right). **b**, Heat map showing the expression of the top thirty marker genes for each significant cell cluster. **c**, Distribution graph of each cell cluster from young and old group. **d**, Dot plot showing expression of marker genes for endothelial subtypes including Fenestrated (FEC), Arterial (Art), and Venous (Ven) endothelial cells. Representative marker genes categorized by endothelial subtypes: pan-endothelial (e.g., *Pecam1*, *Cdh5*, *Cd34*, *Eng*), venous (*Slc38a5*, *Tmsb10*, *Ctla2a*, *Ctss*), arterial (*Gkn3*, *Mgp*, *Sema3g*, *Stmn2*), and fenestrated endothelial cell (FEC)-enriched (*Plvap*, *Col15a1*, *Exoc3l2*, *Esm1*). Dot size reflects the proportion of expressing cells, and color indicates scaled average expression.

**Extended Fig. 3.**
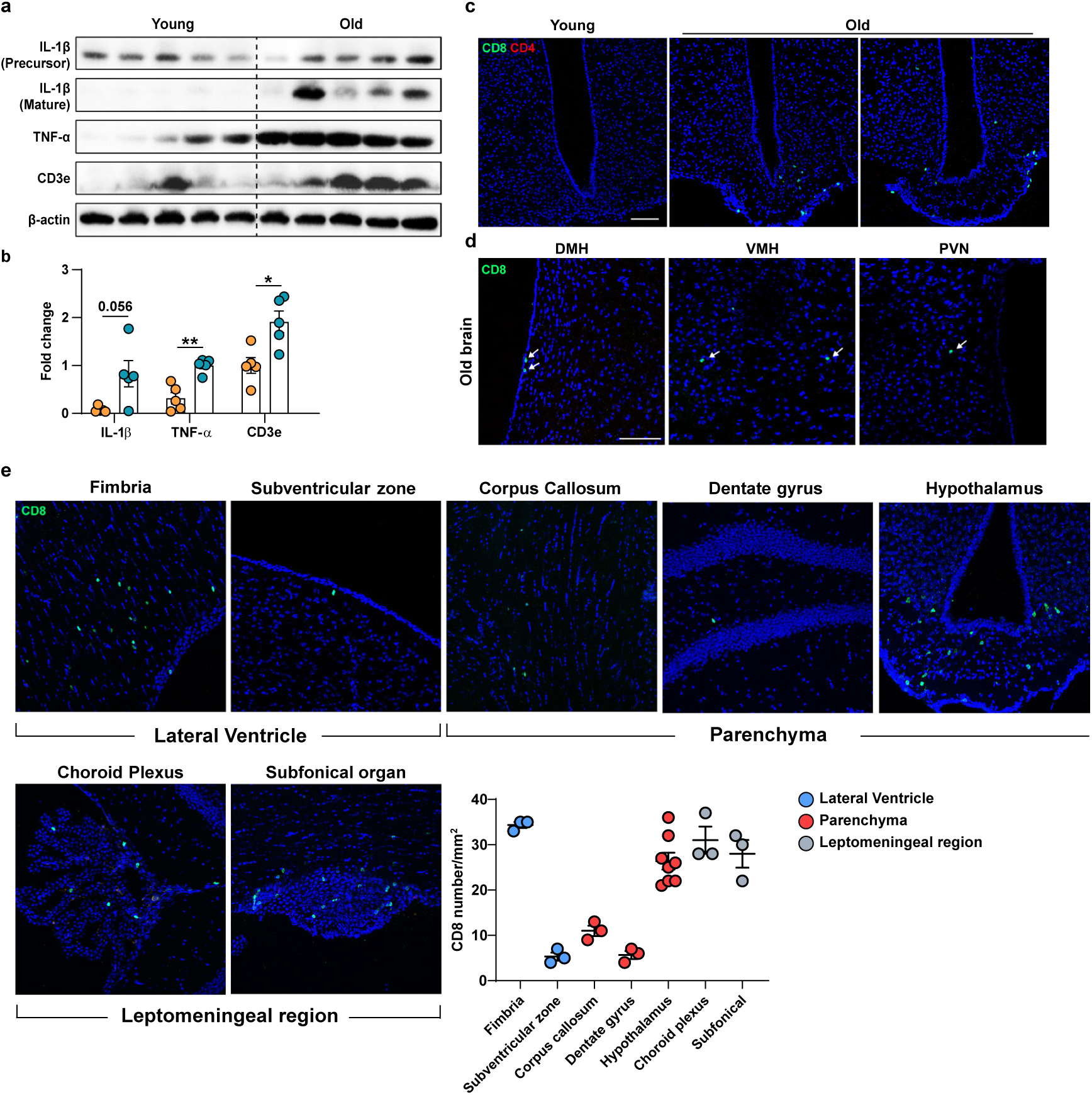
Age-related hypothalamic inflammation and distribution of CD8 infiltration across brain regions. **a**, Immunoblot analysis of precursor and mature IL-1β, TNF-α, and CD3e in hypothalamic lysates from young and old mice. **b**, Quantification of immunoblot band intensities normalized to β-actin (n = 5 per group). **c-d**, Representative immunofluorescence images of CD8 or CD4 in the hypothalamus of young and aged mice (c) and hypothalamic subregion: Dorsomedial hypothalamus (DMH), ventromedial hypothalamus (VMH), Paraventricular nucleus (PVN) from aged mouse brain (d). Scale bar, 100 μm. **e**, Distribution of CD8 across multiple brain regions in aged mice. Quantification of CD8 number per mm² is shown (right) (n = 3 for all regions, except hypothalamus n = 8). Scale bar, 100 μm. Dot plots show mean ± s.e.m. Statistical significance was determined by two-tailed Student’s t-test. *P < 0.05, **P < 0.01.

**Extended Fig. 4.**
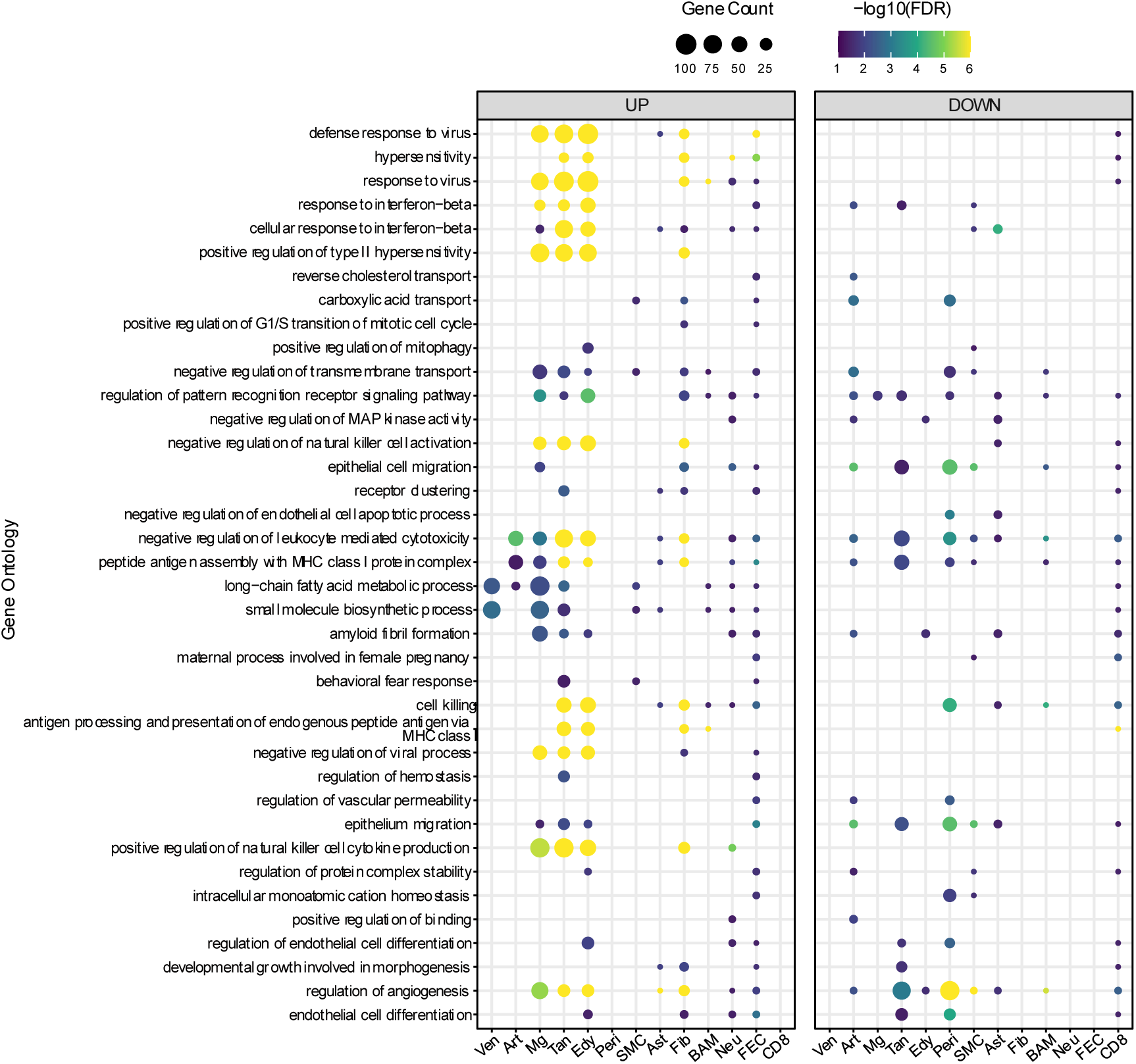
Gene Ontology enrichment for upregulated and downregulated differentially expressed genes (DEGs) from old versus young in hypothalamus. Dot plot indicating Gene ontology (GO) analysis for upregulated and downregulated DEGs of 18 months versus 3 months per cell cluster. Bar color indicates enrichment (–log10(FDR)); dot size indicates the number of marker genes per GO term.

**Extended Fig. 5.**
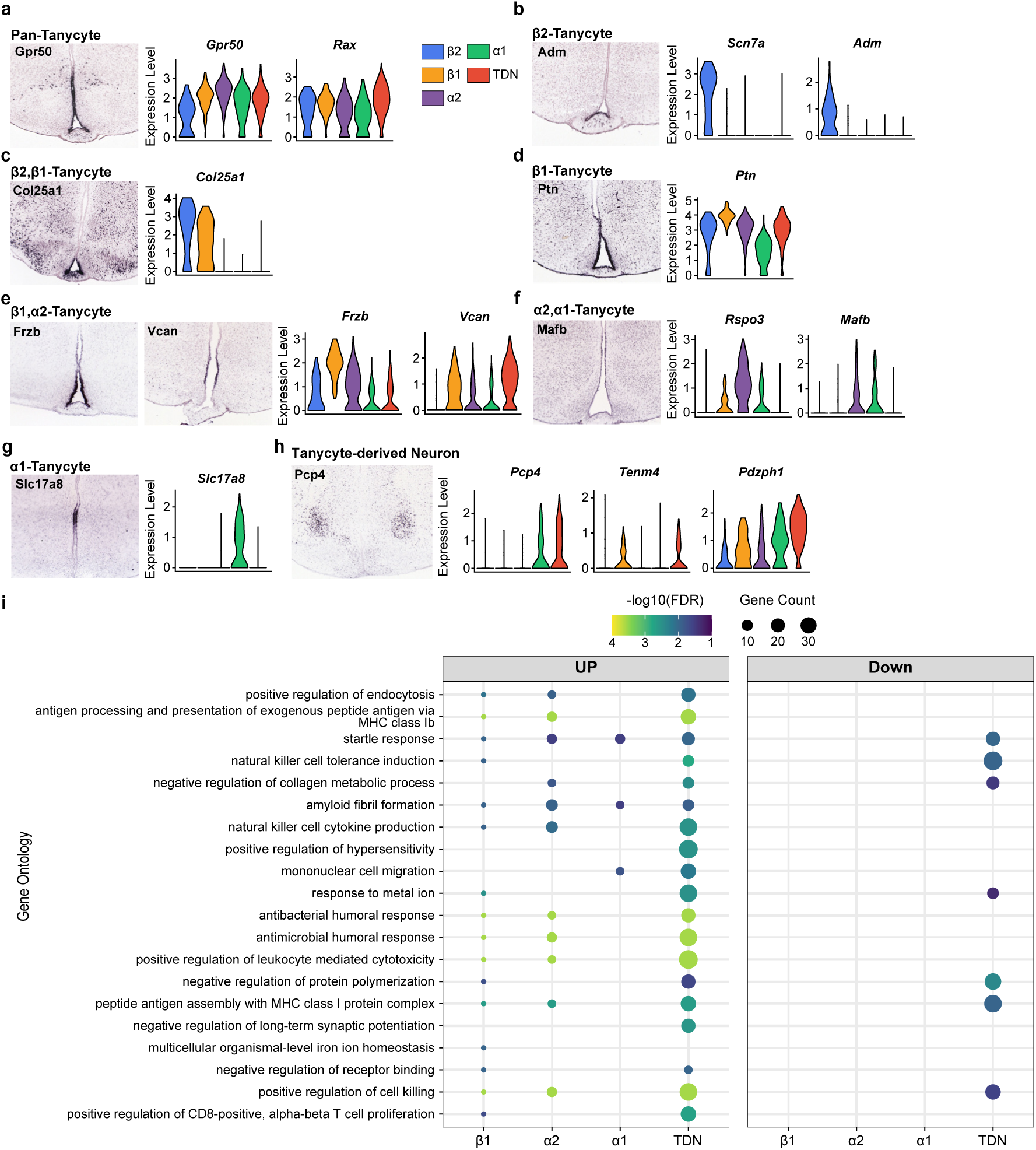
Subclustering analysis of tanycyte from scRNA-seq data. **a-h**, Violin plots and in situ hybridization (ISH) images from Allen Mouse Brain Atlas showing the expression of representative marker genes for each tanycyte subtype—pan-tanycyte (*Gpr50*, *Rax*) (a), β2-tanycyte (*Scn7a*, *Adm*) (b), β2,β1-tanycyte (*Col25a1*) (c), β1^-^ tanycyte (*Ptn*) (d), β1,α2-tanycyte (*Frzb, Vcan*) (e), α2,α1-tanycyte (*Rspo3, Mafb*) (f), α1-tanycyte (*Slc17a8*) (g) and tanycyte-derived neurons (TDN; *Pcp4*, *Tenm4*, *Pdzph1*) (h), based on integrated scRNA-seq profiles from young and aged mice. **i**, Gene Ontology enrichment of DEGs with aging within each tanycyte subtype. The left panel shows terms enriched among genes upregulated with age; the right panel shows downregulated terms. Dot size indicates gene count and color denotes enrichment (−log10(FDR)).

**Extended Fig. 6.**
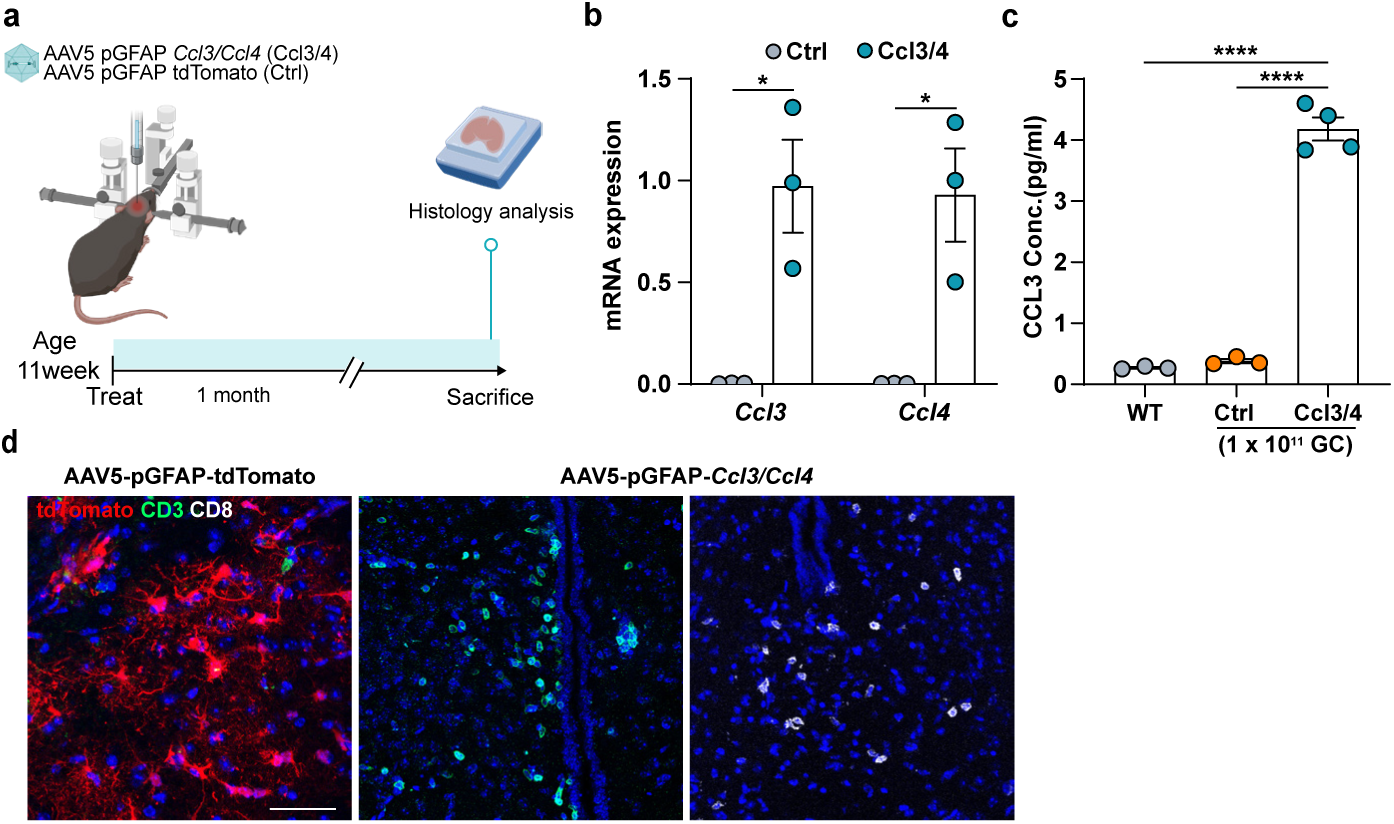
CCL3/4 overexpression induces T cell infiltration in the hypothalamus. **a**, Experimental timeline for stereotaxic injection of AAV5-pGFAP-tdTomato (Ctrl) or AAV5-pGFAP-*Ccl3/Ccl4* (*Ccl3/4*) into the hypothalamus of 11-week-old mice, followed by histological analysis 1 month later. **b**, qRT**-**PCR analysis of hypothalamic tissue confirmed robust overexpression of *Ccl3* and *Ccl4* in the *Ccl3/4*-injected group compared with Ctrl (n = 3 per group). **c**, ELISA measurements showing increased CCL3 protein levels in hypothalamic lysates from Ccl3/4-injected mice compared to WT or Ctrl groups (n = 3∼4 per group). **d**, Representative immunofluorescence images of hypothalamus from mice injected with Ctrl Ccl3/4 AAV, showing CD3 and CD8 infiltration. Scale bar, 50 μm. Dot plots show mean ± s.e.m. Statistical analyses: two-tailed Student’s t-test (**b**) and one-way ANOVA with Tukey’s post hoc test (**c**). *P < 0.05, ****P < 0.0001.

**Extended Fig. 7.**
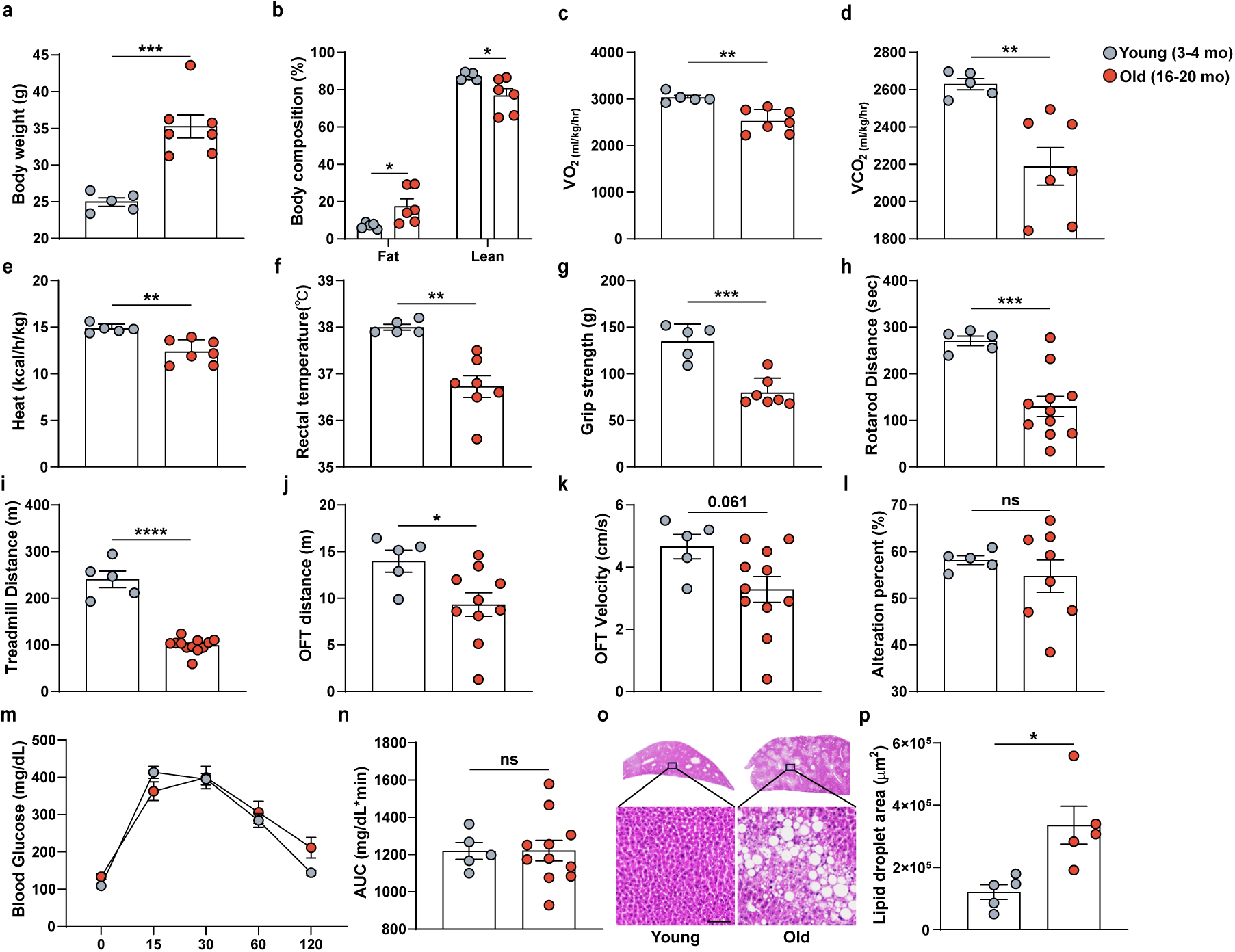
Age-associated changes in systemic metabolism and motor performance. Body phenotype and functional performance in young (3–4 months, n = 5) and old (16–20 months, n = 5∼11) mice: body weight (a), body composition analysis (b), oxygen consumption (vO₂) (c), carbon dioxide production (vCO₂) (d), energy expenditure (e), recta temperature (f), grip strength (g), rotarod performance (h), treadmill performance (i), locomotion distance and velocity in the open field test (j, k), Y-maze (l), glucose tolerance and AUC quantification (m, n), and representative H&E-stained liver sections (o) with quantification of lipid droplet area per field (µm²) (p). Scale bars, 100 µm. Data are shown as mean ± s.e.m.; two-tailed unpaired Student’s t-test. P < 0.05, P < 0.01, *P < 0.001.

**Extended Fig. 8.**
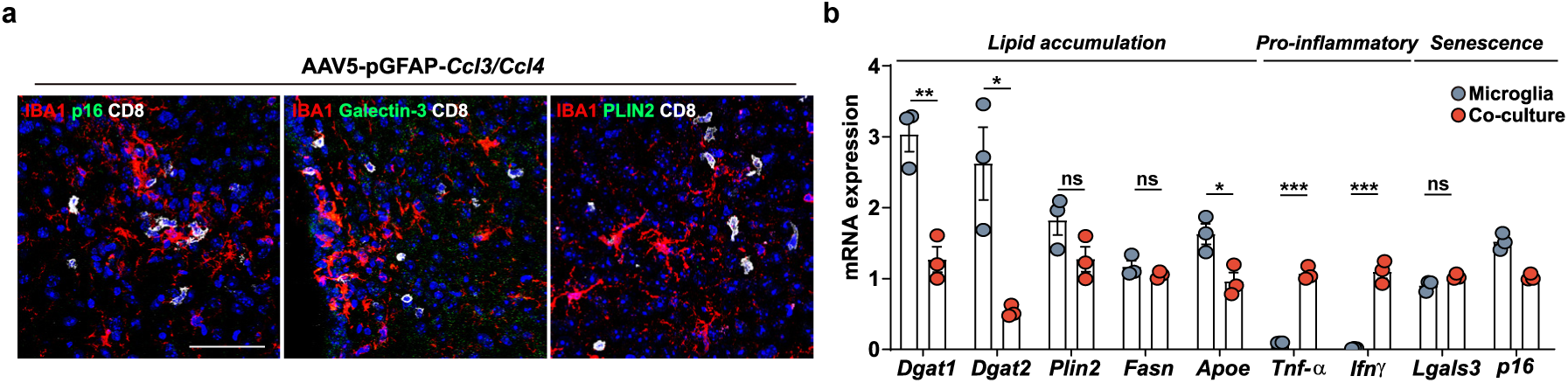
CD8 are not associated with lipid accumulated microglia. **a**, Representative immunofluorescence images of CD8 and IBA1⁺ microglia expressing p16 or Galectin-3, or PLIN2 in hypothalamic sections from young mice injected with AAV5-pGFAP-*Ccl3/Ccl4*. Scale bar, 50 µm. **b**, qRT–PCR analysis of primary microglia cultured alone (Microglia) or co-cultured with activated T cells (Co-culture), examining transcripts associated with lipid accumulation (*Dgat1, Dgat2, Plin2, Fasn, Apoe*), inflammation (*Tnf-α, Ifn-γ*), and senescence (*Lgals3, p16*) (n = 3 per group). Two-tailed unpaired Student’s *t*-test. *P < 0.05, **P < 0.01, ***P < 0.001.

**Extended Fig. 9.**
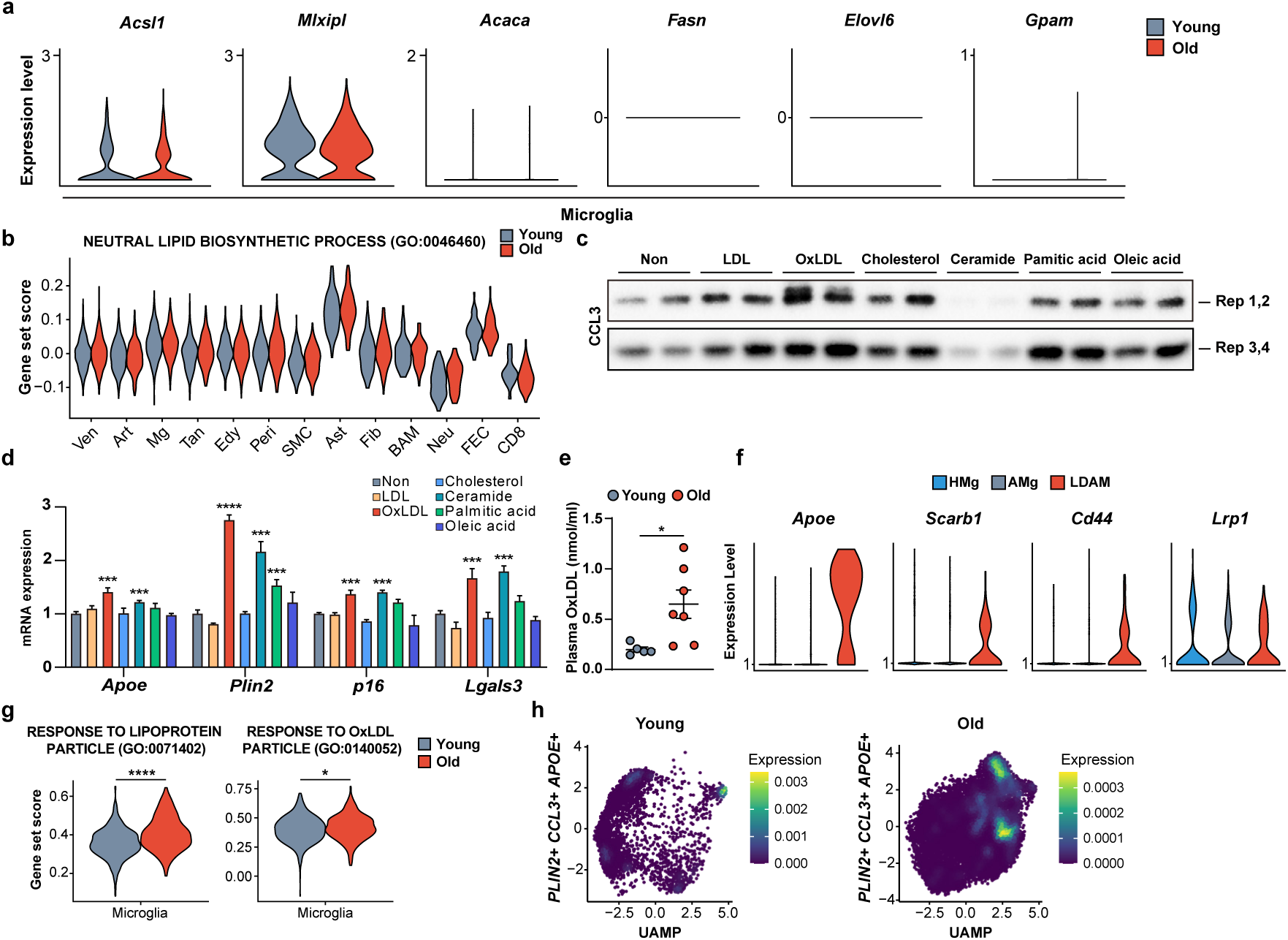
Extrinsic OxLDL exposure induces transformation of hypothalamic microglia into the LDAM subtype. **a**, Violin plots showing age-dependent expression of lipid biosynthesis genes (*Acsl1*, *Mlxipl*, *Acaca*, *Fasn*, *Elovl6*, *Gpam*) in microglia from scRNA-seq datasets. **b**, Module score for GO term “Neutral lipid biosynthetic process” (GO:0046460) across major hypothalamic cell types, comparing young and old mice. **c-d**, BV2 microglia were treated with various lipid species (LDL, OxLDL, cholesterol, ceramide, palmitic acid, oleic acid) to assess their potential to induce CCL3 expression and LDAM-related gene markers. **c**, Immunoblot shows CCL3 protein levels in the conditioned media (n = 4 per group). **d**, qRT-PCR quantification of *Apoe*, *Plin2*, *p16*, and *Lgals3* (n = 3∼6 per group). **e**, ELISA detection for OxLDL-associated protein level from young and aged plasma sample (young, n = 5; old, n =7). **f**, Violin plot showing expression level of scavenger receptor genes (*Apoe*, *Scarb1*, *Cd44*, *Lrp1*) from microglial subcluster of scRNA-seq data. **g**, Module scores for GO term “Response to lipoprotein particle” (GO:0071402) and “Response to oxidized LDL particle” (GO:0140052) in microglia **h**, Kernel density plots showing triple-positive microglia (*PLIN2⁺ CCL3⁺ APOE⁺*) from human HYPOMAP snRNA-seq dataset of young and old human hypothalamus. Dot plots show mean ± s.e.m; two-tailed unpaired Student’s *t*-test. *P < 0.05, ***P < 0.001, ****P < 0.

**Extended Fig. 10.**
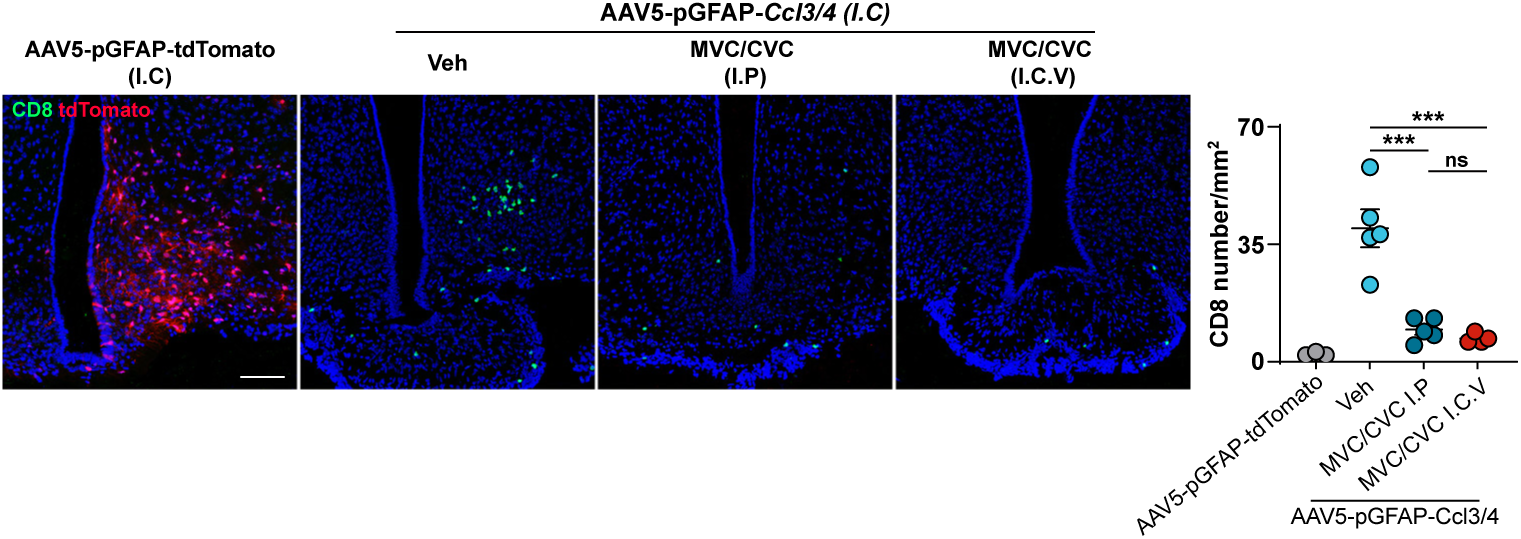
Combined MVC/CVC treatment by intraperitoneal injection blocks intrahypothalamic T cell infiltration. Mice delivered AAV5-pGFAP-*Ccl3/4* by intracranially (I.C) received MVC/CVC or vehicle either intraperitoneally (I.P) or intracerebroventricularly (I.C.V, via osmotic pump) for 2 weeks. Hypothalamic CD8 were quantified (n = 3∼5 per group). Statistical significance was determined using one-way ANOVA followed by Tukey’s post hoc test. ***P < 0.001.

