## Supplemental File for "Pharmacological intervention targeting neuroimmune axis in the aging hypothalamus prevents age-associated physiological decline"

**Supplementary information**

### Supplementary Figure

**Supplementary Fig. 1 Validation annotation of cell clusters by automated sc-type classification.**

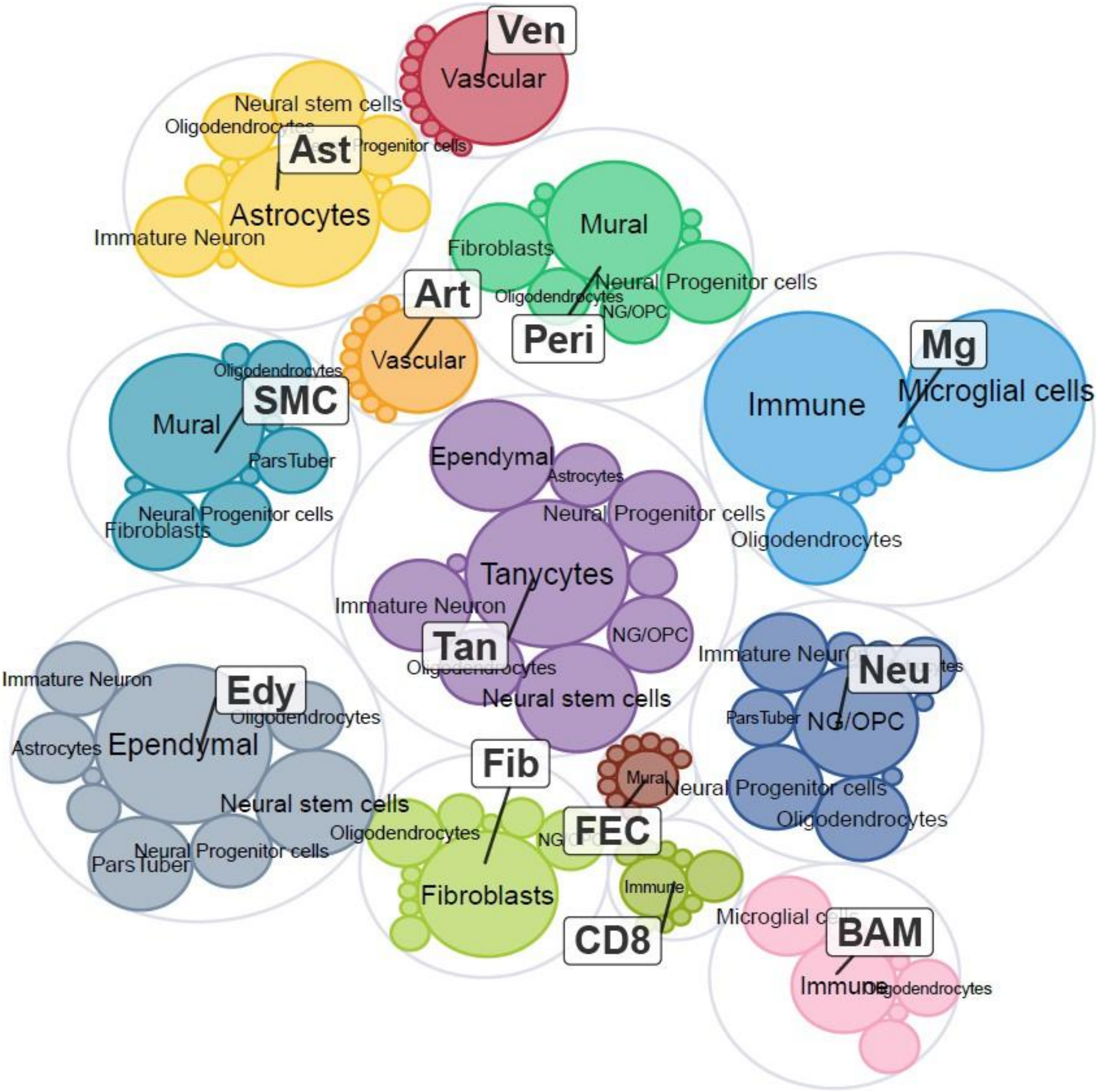

Cell cluster annotation based on sc-type implementation.

**Supplementary Fig. 2 Gene Ontology enrichment of cluster-specific marker genes validates cell-type annotations.**

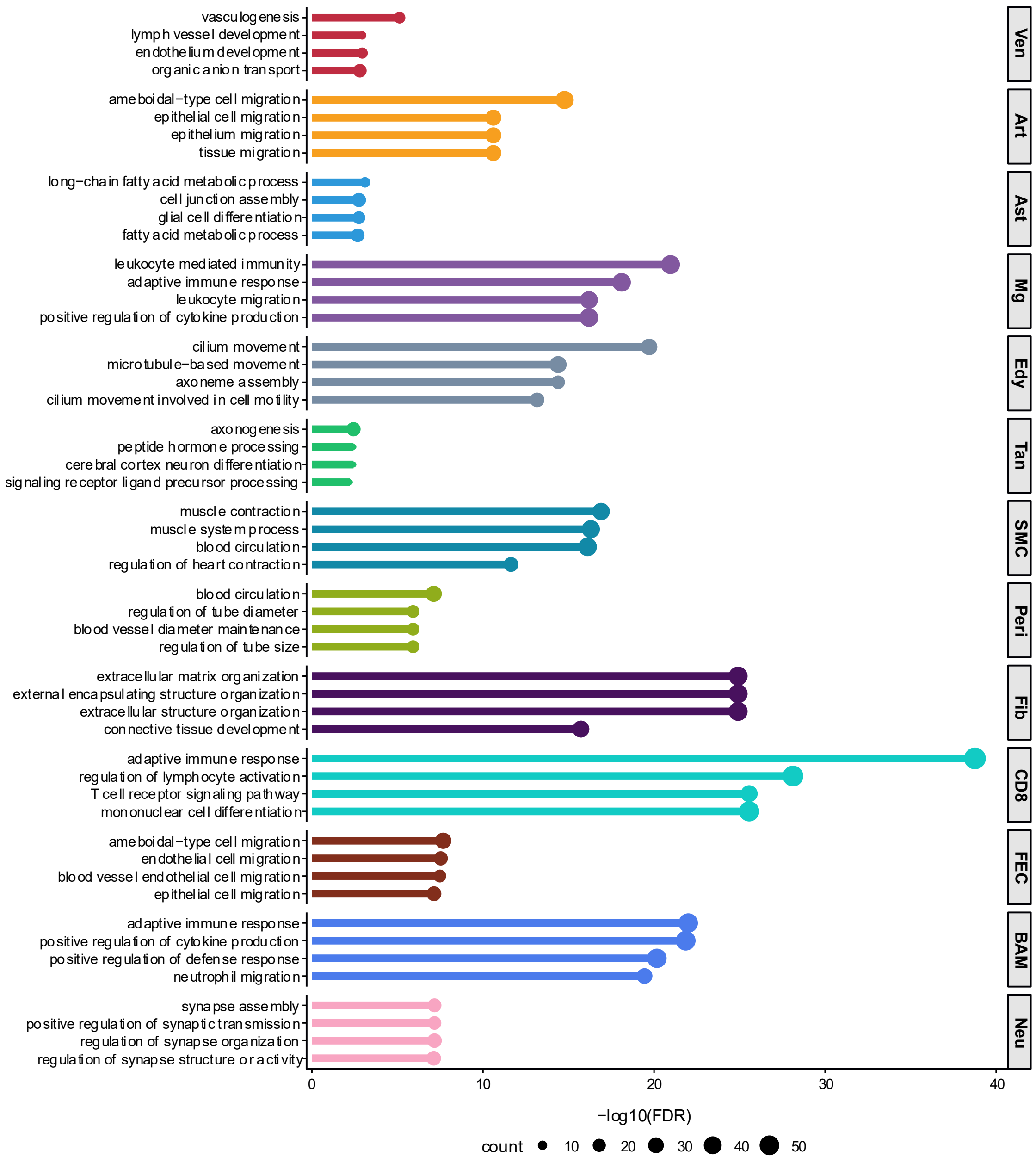

Gene ontology terms enriched for the marker genes of each scRNA-seq cluster from mouse hypothalamus (young and old group combined). Bar length indicates enrichment (–log10(FDR)); dot size indicates the number of marker genes per GO term.

**Supplementary Fig. 3 Phenotypic identity of hypothalamic neural stem cell (htNSC) cultures.**

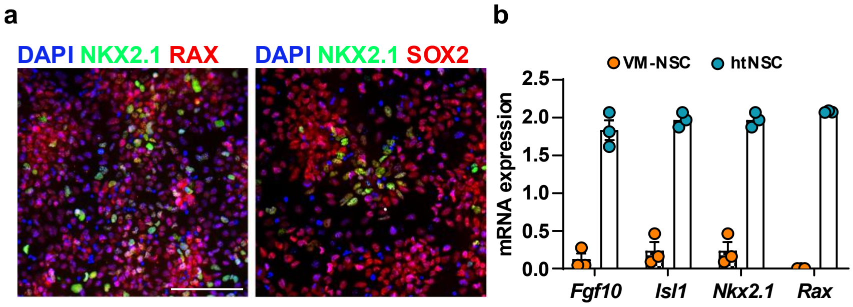

**a,** Representative immunostaining images of htNSCs in culture, showing expression of tanycytic-specific markers (NKX2.1, RAX) and the general NSC marker SOX2. Scale bar, 100µm.

**b,** Comparative mRNA expression of tanycyte-specific marker genes in cultures derived from rodent hypothalamic tissue (htNSCs) versus ventral midbrain neural stem cells (VM-NSCs). Dot plots show mean ± s.e.m.

**Supplementary Fig. 4 Effects of Ccl3/4 overexpression in the middle-aged hypothalamus on systemic body metabolism and physical performance.**

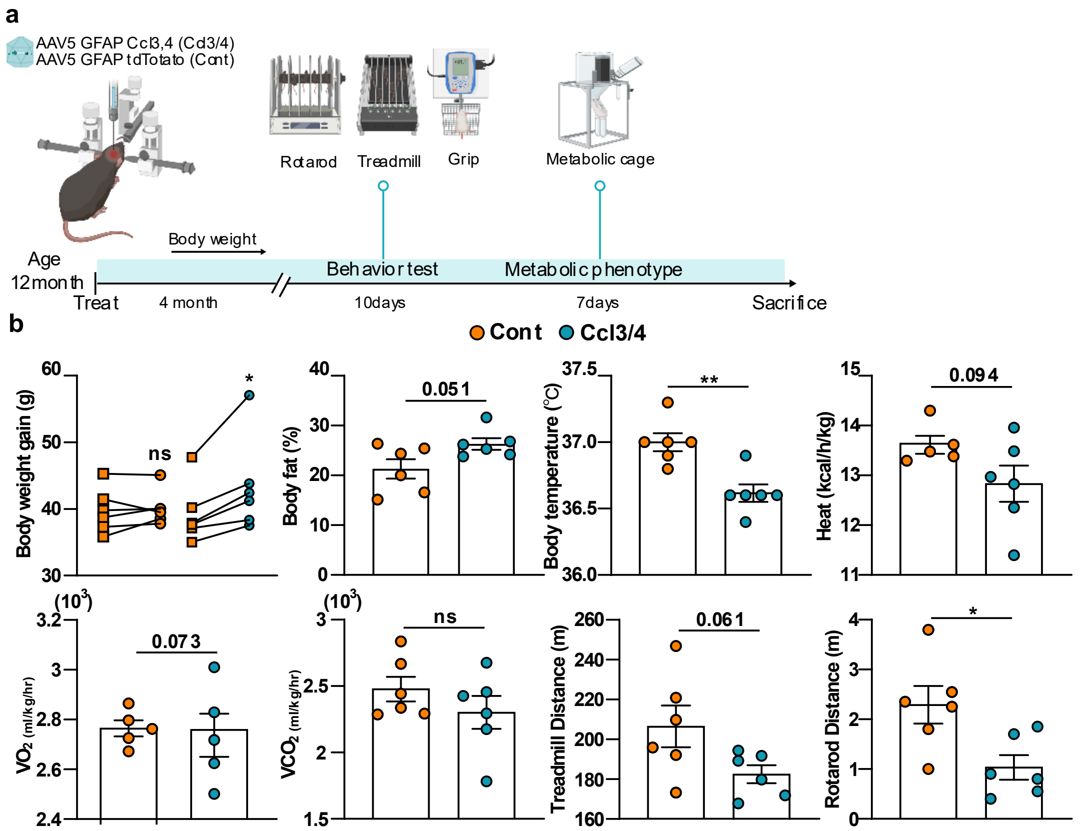

**a,** Experimental timeline for stereotaxic injection of AAV5-GFAP-tdTomato (control) or AAV5-GFAP-Ccl3,Ccl4 (Ccl3/4) into the hypothalamus of 12-month-old mice, followed by metabolic phenotyping and behavioral assessments.

**b,** Body phenotype (body weight gain from pre- to post-injection, fat mass percentage, rectal body temperature) and metabolism (heat production, $VO_2$ and $VCO_2$ consumption), and motor performance (treadmill and rotarod test) assessed in the intrahypothalamic Ccl3/4 expressing and control mice.

Statistical significance was determined by paired t-test for within-group comparison (body weight gain), and unpaired two-tailed Student's t-test for between-group comparisons. *P < 0.05, **P < 0.01.

**Supplementary Fig. 5 Transplanted CD8$^+$ T cell fails to survival in young hypothalamus.**

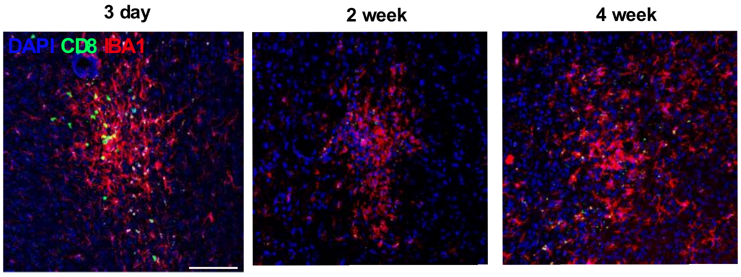

Time-course analysis of CD8$^+$ T cell survival in the hypothalamus following CD8$^+$ T cell transplantation. Immunofluorescence images show survival of CD8$^+$ T cells in injected site with abundant IBA1$^+$ microglial population at 3 days, 2 weeks, and 4 weeks post-surgery. Scale bar, 100µm.

**Supplementary Fig. 6 Hypothalamus-specific enrichment of PLIN2 reflects regional lipid droplet accumulation.**

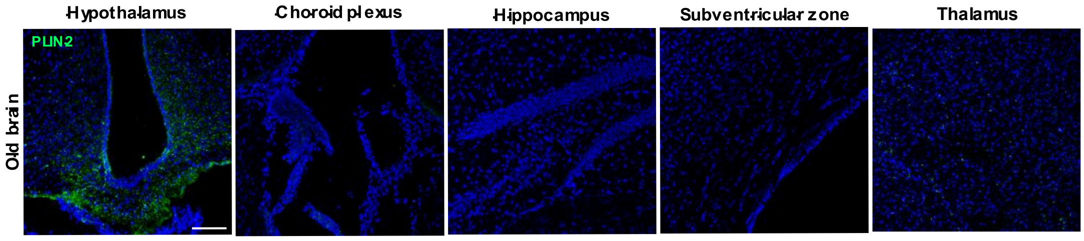

Representative immunofluorescence images of PLIN2 protein across brain regions of aged mice. Scale bar, 100µm.

### Supplementary Methods

Details of reagents, antibodies, primers, apparatus, and equipment used in this study are provided in the Supplementary Lists.

#### Cell culture

*Hypothalamic Neural Stem Cell (htNSC)*

Neural progenitor cells (NPCs) with the potential to differentiate into tanycyte-like NPCs were isolated from the hypothalamic region of embryonic day 12.5 (E12.5) mouse embryos. Dissected hypothalamic tissues were gently dissociated in Hank's Balanced Salt Solution (HBSS) using mechanical trituration. The resulting single-cell suspension was plated onto poly-L-ornithine/fibronectin-coated culture plates.

Cells were maintained in a modified N2 medium supplemented with 64 µg/L insulin, 1% penicillin-streptomycin, 20 ng/mL recombinant human basic fibroblast growth factor (bFGF), and B-27 supplement without vitamin A. NPCs were allowed to expand for 48 hours prior to passaging for downstream experiments. To validate the identity and regional specificity of cultured hypothalamic NPCs, immunostaining was performed for key hypothalamic tanycytic NPC markers, including SOX2, RAX, and NKX2.1.

*HEK 293T Cell line*

Human embryonic kidney 293T (HEK 293T) cells were maintained in 100-mm dishes at 37 °C and 5% $CO_2$. Cells were cultured in Dulbecco's Modified Eagle Medium (DMEM) supplemented with 10% fetal bovine serum (FBS) and 1% penicillin-streptomycin (Pen-Strep). Medium was changed every 2 days, and cells were split every 2–3 days at 70–80% confluency using 0.05% trypsin-EDTA. For transfection, cells were seeded to reach 70–80% confluency within 24 hours. Only cells that had undergone at least 2–3 passages post-thaw were used for experiments. All cultures were routinely tested for mycoplasma contamination, and only mycoplasma-negative cells were used.

*BV2 Microglial Cell Line*

BV2 murine microglial cells were cultured in 100-mm tissue culture dishes at 37 °C in a humidified atmosphere containing 5% $CO_2$. Cells were maintained in DMEM supplemented with 10% FBS, 1% Pen-Strep and 1% GlutaMAX. Culture medium was refreshed every 2 days, and cells were passaged every 3–4 days using 0.05% trypsin-EDTA when cultures reached ~80% confluency.

For plasmas supplementation and component treatment to BV2 cells, cells were seeded at a density of $3 \times 10^5$ cells per well in 6-well plates or $4 \times 10^4$ cells per well in 24-well plates. After 24 hours, the medium was replaced with serum-free DMEM containing 1% Pen-Strep and 1% GlutaMAX. The following reagents were then added to each well and incubated for 24 hours: oxidized low-density lipoprotein (OxLDL, 50µg/mL), ceramide (20µM), palmitic acid (50µM), Oleic acid (50µM), low-density lipoprotein (LDL, 50µg/mL), and cholesterol (25µg/mL). After treatment, cells and conditioned media (CM) were immediately harvested for subsequent analyses.

*Primary Microglia*

Primary microglia were isolated from postnatal day 1 (P1) ICR mouse pups. Whole brains were dissected and dissociated, and mixed glial cultures were established in T75 flasks using DMEM/F12 medium supplemented with 10% FBS, 10% horse serum, 1% GlutaMAX, B27 supplement, 1% Pen-Strep, 1mg/mL D-glucose, and 10ng/mL bFGF. Cultures were maintained for 10 days with media changes every 2 days. For microglial isolation, flasks were shaken on an orbital shaker at 220 rpm for 2 hours at 37 °C. The supernatant containing detached microglia was collected, centrifuged at 300 g for 5 minutes, and replated onto 6-cm dishes for further analysis.

*$CD8^+$ T cell Isolation and Coculture*

$CD8^+$ T cells were isolated from mouse spleens using magnetic-activated cell sorting (MACS). Briefly, spleens were mechanically dissociated in HBSS, filtered through a 70-µm strainer, and centrifuged at 300 × g for 3 minutes. Red blood cells were lysed with ACK buffer, and splenocytes were resuspended in MACS buffer (0.5% BSA, 2 mM EDTA in PBS). $CD8a^+$ T cells were labeled with biotin-conjugated antibody and anti-biotin microbeads from isolation kit and cells were passed through LS columns to obtain a negatively selected $CD8^+$ T cell population. Enriched cells were resuspended in T cell culture medium (RPMI-1640 supplemented with 10% FBS, 1% Pen-Strep, 50µM β-mercaptoethanol and 5ng/ml IL-2). To activate and expand $CD8^+$ T cells, isolated cells were cultured for 5 days using Dynabeads™ Mouse T-Activator CD3/CD28 for T-Cell Expansion and Activation, following the manufacturer's protocol.

For co-culture experiments activated $CD8^+$ T cells were seeded at a 1:5 ratio (T cell: htNSC) on 6 well plate with either primary microglia or hypothalamic neural stem cell. Co-cultures were maintained for 48 hours in appropriate media without additional cytokine supplementation. Following incubation, cells were collected for downstream assays including flow cytometry, qPCR, and Western blotting or imaging analyses.

**$CD8^+$ T cell Migration Assay**

To evaluate the chemotactic response of $CD8^+$ T cells to factors secreted by OxLDL-treated microglia, a transwell migration assay was performed using 24-well transwell plates equipped with 5.0-µm pore size inserts. The lower chamber was filled with 500µl of conditioned medium collected from OxLDL-treated BV2 microglia or control medium. A total of 4,000 purified $CD8^+$ T cells, suspended in 100µl of medium, were seeded into the upper chamber. Following a 4 hours incubation at 37 °C in a humidified atmosphere containing 5% $CO_2$, cells that had migrated to the lower chamber were collected and counted using a hemocytometer. The migration efficiency was calculated as the percentage of cells that migrated relative to the initial number of cells seeded in the upper chamber.

**htNSC Cytotoxicity Assay in a Transwell System**

Hypothalamic neural stem cells (htNSCs) were seeded in the lower wells of 24-well transwell (5.0-µm pore inserts) and maintained in T-cell medium supplemented with BV2 conditioned medium (OxLDL–treated or untreated). Activated $CD8^+$ T cells were added to the upper inserts and exposed to maraviroc (10µM) and cenicriviroc (10µM) or vehicle for the duration of co-culture. After incubation for 48 hours at 37 °C in 5% $CO_2$, inserts were removed and htNSC death was quantified using ethidium homodimer-1 (EthD-1) staining.

**Oil Red O Staining**

To assess intracellular lipid accumulation, cultured cells were fixed with 4% paraformaldehyde (PFA) for 15 minutes at room temperature and rinsed twice with phosphate-buffered saline (PBS). Fixed cells were incubated with 100% propylene glycol for 5 minutes to enhance dye permeability. Subsequently, cells were stained with pre-warmed Oil Red O solution (60°C) for 30 minutes. After staining, cells were briefly differentiated in 85% propylene glycol for 1 minute, followed by two washes with distilled water. Nuclei were counterstained with hematoxylin. Coverslips were mounted using an aqueous mounting medium and bright imaged using a Leica DM5000B microscope.

**Mouse Tissue Isolation and Histology Analysis**

Mice were deeply anesthetized, and blood was collected via transcardial perfusion. Subsequently, major organs were dissected, including the brain and liver. For immunohistochemical analysis, brain and liver tissues were post-fixed in 4% PFA at 4 °C overnight, followed by cryoprotection in 30% sucrose for 48 hours. To assess hepatic lipid accumulation, liver sections were subjected to hematoxylin and eosin (H&E) staining. Images were acquired using a AxioScan.Z1 and analyzed using ImageJ software for immunoblotting and ELISA analysis, dissected tissue is directly moved on solution and lysed for downstream process.

**Preparation of Single-Cell Suspensions from Mouse Hypothalamus**

Hypothalamic tissue dissociation was performed using Multi tissue dissociation kit 1 and gentleMACS Dissociator according to manufacturer's instruction. After digestion, cell suspensions were filtered through a 70-µm cell strainer to remove undissociated clumps. Myelin debris was removed using Debris Removal Beads to enrich viable single cells. Cell viability and concentration were assessed using PI staining and cell counting. Only single-cell suspensions with >85% viability were used for downstream single-cell RNA sequencing.

**Single-Cell RNA Sequencing**

Single-cell RNA-seq libraries were prepared using the Chromium Next GEM Single Cell 5′ and V(D)J Library Kits (10x Genomics) following the manufacturer's instructions. Briefly, single-cell suspensions were adjusted to a target input of 10,000 cells and loaded onto a Chromium Next GEM Chip K with Gel Beads and Partitioning Oil to generate Gel Beads-in-Emulsion (GEMs). Within each GEM, RNA transcripts were reverse-transcribed and arcoded with cell- and molecule-specific barcodes.

For 5′ gene expression libraries, full-length cDNA was amplified, end-repaired, A-tailed, and ligated with sequencing adapters. PCR enrichment was then performed to produce sequencing-ready libraries. For V(D)J libraries, cDNA containing T cell receptor (TCR) or B cell receptor (BCR) transcripts were selectively amplified using specific primer mixes (T Cell Mix1/2) to capture full-length variable regions.

The libraries were sequenced on an Illumina HiSeq platform following the manufacturer's protocol. Sequencing reads were processed using the Cell Ranger pipeline (10x Genomics). Reads were aligned to the mouse reference genome (mm10-2020-A, GRCm38) and assembled per cell based on Unique Molecular Identifiers (UMIs) and 10x barcodes to generate high-confidence gene expression matrices and TCR/BCR clonotype data for downstream analysis.

*Pre-processing and Quality Control of scRNA-seq Data*

Gene expression matrix was cleaned using Cellbender (v 0.3.0) remove background command with expected-cells 9000 argument. The resulting h5 files were then loaded into R (v4.4.2) using helper function Read_CellBender_h5_Mat from scCustomize (v.3.0.1). Cells were filtered using the following thresholds: 3,000 ≤ nCount ≤ 30,000, 2,000 ≤ nFeature ≤ 8,000, percent.mito < 5%, percent.ribo >2%. Doublet was removed using scDblFinder (v1.18.0).

*Data Integration, Dimensionality Reduction, and Clustering*

For each sample, a Seurat object was created (min.cells = 10, min.features = 5). The samples were then integrated using "HarmonyIntegration". NN and SNN graph were constructed using FindNeighbors function using the first 35 dimensions. UMAP were performed using "umap-learn" method with 200 learning epochs. FindClusters (resolution = 0.28) was performed using Louvain algorithm with multilevel refinement (algorithm =2). This setting was chosen to balance the number of cluster and biologically meaningful separation. Cell type annotation was performed manually using established cell marker: Tmsb10 (Vein), Gkn3 (Artery), Agt (Astrocyte), Cx3cr1(Microglia), Ccdc153 (ependymal cells), Gpr50 (tanycytes), Acta2 (Smooth muscle cell), Kcnj8 (Pericyte), Tcf21 (Fibroblast), Cd8a ($CD8^{+}$ T cells), Plvap (Fenestrated endothelial cell), Pf4 (border-associated macrophages), and Snap25 (Neuron). Cell type annotation was validated using sc-type with the brain reference from ScTypeDB with slight modification to add tanycyte marker. FindAllMarkers was implemented to find top genes for each cluster and top 200 genes for each cluster was used for GO analysis to further confirm each cluster identity based on the functional ontology.

*Differentially Expressed Genes and Gene Ontology Analysis*

Differential gene expressions between age groups for each cluster were performed using the FindMarkers() function in Seurat with default parameters. Genes with an adjusted *P*-adjusted value < 0.05 and absolute $\log_2$ fold change > 0.585 (1.5 fold) were considered significant. GO BP enrichment analysis was performed using the clusterProfiler package (v4.12.1) based on implementation Org.Mm.eg.db (v.3.19.1). Redundant term was semantically reduced with GOSemSim and rrvgo package. Results were shown as reduced dot plots on a global GO map arranged hierarchically based on semantic distance.

*TCR Sequencing and Functional CD8 Analysis*

10x V(D)J outputs (TRA/TRB) summarized with scRepertoire (v2.0.7); clonotype categories (iNKT/MAIT/conventional) and clonal dominance distributions compared by age. Functional state of CD8 was estimated by projecting CD8 cluster onto the "mouse TIL" reference using ProjecTILs. State composition was predicted using plot.statepred.composition function.

*Human HYPOMAP Microglia Subclustering*

HYPOMAP RDS file was obtained from the original publication[1]. Only microglia ("C2-49") subset was analyzed in our study. One donor (DonorID = 3u5kk) was excluded due to excessive ambient oligodendrocyte RNA contamination. The donor was stratified into two groups: young (≤65 years) and old (>65 years). NN and SNN graph construction was performed using all of the dimensions in the existing scvi harmonization. Due to the high sparsity of snRNAseq data, gene expression was visualized using the Nebulosa (v1.14.0) package.

**Bulk RNA-seq Transcriptome Analysis**

Total RNA was isolated from hypothalamic tissues of mice at 7 weeks, 12 months, and 24 months of age using TRIzol reagent. RNA concentration was measured using the Quant-iT RiboGreen RNA assay kit, and RNA integrity was assessed with the TapeStation RNA ScreenTape system. Only samples with RNA integrity number (RIN) > 7.0 were used for library preparation.

For each sample, 1 μg of high-quality total RNA was used to construct sequencing libraries using the TruSeq Stranded mRNA Sample Prep Kit. Library quality and size distribution were validated using the TapeStation D1000 ScreenTape (Agilent, #5067-5582), and library quantification was performed with the KAPA Library Quantification Kit for Illumina platforms (KAPA Biosystems, #KK4854) according to the manufacturer's protocol. Indexed libraries were sequenced on the Illumina NovaSeq 6000 platform at Macrogen (Seoul, South Korea).

Raw sequencing reads were aligned to the mouse reference genome (mm10-2020-A, GRCm38) using HISAT2 (v2.1.0). Transcript- and gene-level abundances were quantified as read counts and fragments per kilobase of transcript per million mapped reads (FPKM). Differential gene expression analysis was conducted using the DESeq2 R package (1.46.0), and significantly differentially expressed genes (DEGs) were defined by an adjusted p-value $< 0.01$ and $|\log2 \text{ fold change}| > 1$. Gene Ontology (GO) enrichment analysis of DEGs was performed using the clusterProfiler R package (v4.12.1). Data visualization, including volcano plots and GO enrichment bar graphs, was generated using ggplot2 (3.5.2), and heatmaps were created with the pheatmap function in R.

**T cell Staining in Postmortem Human Hypothalamus Tissue Sample**

Formalin-fixed paraffin-embedded (FFPE) human hypothalamus brain tissues were sectioned and mounted on glass slides. Sections were deparaffinized in xylene, rehydrated through a graded ethanol series, and subjected to antigen retrieval in citrate buffer (pH 6.0) at 95–100 °C for 25 minutes. After cooling to room temperature, tissues were permeabilized with 0.3% Triton X-100 in PBS for 10 minutes and incubated in 3% hydrogen peroxide in distilled water for 10 minutes to quench endogenous peroxidase activity.

Endogenous biotin and avidin binding sites were blocked using an Avidin/Biotin Blocking Kit, followed by incubation in blocking buffer (1% BSA 10% normal donkey serum in PBS) for 1 hour at room temperature. Sections were then incubated overnight at 4 °C with primary antibodies against CD3e and CD8, diluted in blocking buffer. Next, sections were incubated with the appropriate biotinylated secondary antibodies (Goat anti-rabbit-biotinylated) for 1 hour at room temperature, followed by ABC reagent and streptavidin–HRP for 30 minutes. Colorimetric detection was performed using a DAB Substrate Kit, and sections were counterstained with hematoxylin, dehydrated in graded ethanol, cleared in xylene, and coverslipped with permanent mounting medium.

Brightfield images were acquired using a Leica DM5000B microscope. For each sample, the hypothalamic ventricular zone was imaged at 20x or 40x magnification.

**Immunohistochemistry for Mouse Brain Sections and Immunocytochemistry for in vitro Cells**

Following post-fixation, mouse brains were dehydrated in 30% sucrose for 48 hours, embedded in optimal cutting temperature (OCT) compound, and coronally sectioned at a thickness of 30µm. Sections were mounted on glass slides and rinsed with PBS. Antigen retrieval was performed using citrate buffer (pH 6.0) at 80 °C for 15 minutes. Blocking was performed in PBS containing 5% normal donkey serum, 1% BSA, and 0.2% Triton X-100 for 1 hour at room temperature.

Primary antibody staining was carried out overnight at 4 °C using the same blocking buffer. After incubation, sections were washed with PBS, incubated with secondary antibodies for 1 hour at room temperature, and washed again. To reduce tissue autofluorescence, sections were treated with TrueBlack for 1 minute. Slides were mounted using VECTASHIELD Antifade Mounting Medium with DAPI and stored at 4 °C until imaging.

For immunocytochemistry (ICC) of cultured cells, cells were fixed in 4% PFA for 15 minutes at room temperature, rinsed in PBS, and blocked/permeabilized in PBS containing 1% BSA and 0.2% Triton X-100 for 1 hour. Primary antibody incubation was performed overnight at 4 °C, followed by PBS washes and incubation with appropriate fluorescent secondary antibodies for 1 hour at room temperature. Nuclei were counterstained with DAPI, and coverslips were mounted with antifade mounting medium. For detecting death cell in vitro cultured cell, EthD-1 staining proceeded in live cell before fixation. 2µM EthD-1 solution were added into live cell and incubation for 30 minutes followed by fixation. Details of all antibodies used are provided in Supplementary List.

**Confocal Image Quantification**

Coronal brain sections per mouse were used for immunofluorescence staining. Confocal images were acquired using a Stellaris 5 LIA confocal microscope at either 20x or 40× or 60× magnification. For each antibody staining set (e.g., IFITM3/IBA1, B2m/IBA1), identical laser power, detector gain, and Z-stack acquisition settings were applied across all experimental conditions to ensure consistency. Image analysis and quantification were performed using Fiji.

For quantification of $CD8^+$ T-cell populations in human postmortem or mouse hypothalamic tissue, $CD8^+$ cells were manually counted within a predefined hypothalamic region of interest (ROI) on 20× confocal images, and counts were normalized to area. For microglial co-expression analyses, the mediobasal hypothalamus (MBH) was manually delineated and double-positive microglia were enumerated—defined as $IBA1^+$ cells co-expressing Galectin-3, or $IBA1^+$ cells co-expressing one of the target proteins (IFITM3, B2m, p16, CCl3, or PLIN2). Values were averaged per mouse for group comparisons.

For quantification of Ifitm3 fluorescence intensity in tanycytes, the tanycytic layer lining the ventral third ventricle was manually outlined. Two regions of interest (ROIs) per section were selected, and the fluorescence intensity was measured. Background signal was subtracted, and the resulting values were normalized to the ROI area. The mean intensity per mouse was used for statistical analysis.

All quantification steps were performed using consistent thresholding and analysis parameters across samples.

**Flow Cytometry Analysis**

To assess apoptosis in NSCs following co-culture with T cells, NSCs and activated T cells were co-seeded at a 5:1 ratio (50,000 NSCs: 10,000 T cells) per well. Cells were incubated for 48 hours. After incubation, cells were detached by aspirating the medium and adding 100µl of Accutase to each well, followed by incubation at 37 °C for 5 minutes. The resulting cell suspensions were washed with FACS buffer (PBS + 2% FBS) and stained with Annexin V-FITC and Propidium Iodide (PI) using the Annexin V/PI Apoptosis Detection Kit for 15 minutes at room temperature in the dark.

After staining, samples were washed, centrifuged at 300g for 5 minutes, and resuspended in FACS buffer. Flow cytometric analysis was performed on a FACS Canto II, and data was analyzed using FlowJo software (v10.8.1). Cells were gated based on forward and side scatter to exclude debris and doublets. Apoptotic cells were identified as Annexin $V^+$ $PI^-$ (early apoptosis) or Annexin $V^+$ $PI^+$ (late apoptosis). To validate the staining and gating strategy, the following controls were included: (i) unstained cells, (ii) single-stained controls for Annexin V-FITC and PI.

**Body Composition and Metabolic Phenotyping**

Body weight and food intake were measured weekly using a calibrated digital scale. Whole-body composition, including lean mass and fat mass, was assessed using a quantitative nuclear magnetic resonance (NMR)-based body composition analyzer. Mice were briefly placed into the scanning chamber for non-invasive analysis, and total fat and lean tissue mass were automatically quantified.

To evaluate energy expenditure and metabolic activity, mice were individually housed in a metabolic cage system equipped with a metabolic environmental chamber that maintains thermoneutral condition. Mice were acclimated to the single housing cages for 48 hours prior to the start of metabolic measurements, followed by a continuous 48 hours monitoring period under a 12 h light: 12 h dark cycle. Food and water were provided ad libitum during the entire recording period, using the same diet as in the standard housing conditions.

Oxygen consumption ($VO_2$), carbon dioxide production ($VCO_2$), respiratory exchange ratio (RER), locomotor activity, and energy expenditure were recorded and analyzed using the manufacturer's software. All measurements were performed at the Korea Mouse Phenotyping Center (KMPC), Seoul National University.

**Stereotaxic Surgery**

AAV5- GfaABC1D -tdTomato or a combination of AAV5- GfaABC1D -Ccl3 and AAV5- GfaABC1D -Ccl4 (1µL total volume, $1 \times 10^{11}$ genome copies/µL) was bilaterally injected into the mediobasal hypothalamus (coordinates relative to bregma: −1.2 mm posterior, ±0.3 mm lateral, −5.6 mm ventral from the dura) using a stereotaxic frame apparatus. Mice were anesthetized by intraperitoneal (I.P.) injection of a mixture of Zoletil 50 (0.1mg/kg; Virbac, Carros, France) and Rompun (93.28µg/kg; Virbac). A 26-gauge needle was slowly lowered to the target site, and following injection, the needle was left in place for 25–30 minutes to minimize reflux before being slowly withdrawn.

**CCR5 Antagonist Delivery**

For long-term intracerebroventricular (I.C.V.) delivery of CCR5 antagonists, osmotic pumps were implanted according to the manufacturer's instructions and previously published protocols[2,3] .

**Intracerebroventricular Infusion via Osmotic Pump**

For long-term I.C.V. delivery of CCR5 antagonists, osmotic pumps were implanted according to the manufacturer's instructions (Alzet). Briefly, Maraviroc (625µg/mL) and Cenicriviroc

(125μg/mL) were loaded into mini osmotic pumps connected to brain infusion cannulae (Brain Infusion Kit #3, Alzet) and pre-soaked in sterile PBS at 37 °C for ~40 hours under aseptic conditions.

Mice were anesthetized by intraperitoneal injection of a mixture of Zoletil 50 (0.1mg/kg; Virbac, Carros, France) and Rompun (93.28μg/kg; Virbac), and placed in a stereotaxic apparatus. Cannulae were implanted into the lateral ventricles (coordinates relative to bregma: 0.0 mm anteroposterior, +0.9 mm mediolateral, −2.8 mm dorsoventral from the skull surface) and fixed in place using Loctite 454 adhesive (Henkel). Pumps were inserted subcutaneously in the interscapular region, and incisions were closed with braided sutures (SK 434, 18 mm; Ailee) and surgical adhesive. Mice were monitored daily for recovery and general health during the 14-day infusion period.

**Systemic Administration via intraperitoneal Injection**

For peripheral delivery of CCR5 antagonists, aged mice (16 months old) received intraperitoneal injections of Maraviroc (50mg/kg) and Cenicriviroc (10mg/kg) or vehicle control (20% DMSO, 30% PEG300, 5% Tween 80, 45% saline) once every 3 days for a total duration of 2 months.

**ELISA**

CCL3 and OxLDL levels were measured from mouse hypothalamic tissue lysates, plasma, or BV2 cell-conditioned media using the Mouse MIP-1α (CCL3) Uncoated ELISA Kit and Mouse Oxidized LDL ELISA Kit, respectively, according to the manufacturers' instructions. Absorbance was measured using a microplate reader, and concentrations were calculated based on standard curves.

**Western Blotting**

Proteins were extracted from cultured cells, conditioned media, or mouse tissues using RIPA lysis buffer (50 mM Tris-HCl, pH 7.4; 150 mM NaCl; 1% Triton X-100; 1% sodium deoxycholate; 0.1% SDS; 1 mM EDTA) supplemented with protease inhibitor cocktail and phosphatase inhibitor cocktail. For protein isolation from conditioned media, chloroform–methanol precipitation was performed. Lysates were briefly sonicated and centrifuged at 15,000 × g for 5 minutes at 4 °C to remove insoluble debris. Protein concentrations were determined using the Pierce BCA Protein Assay Kit, according to the manufacturer's instructions.

Equal amounts of protein (15–40μg) were separated by SDS–PAGE and transferred to PVDF membranes. Membranes were blocked with 5% bovine serum albumin (BSA) in TBST (Tris-buffered saline with 0.1% Tween-20) for 1 hour at room temperature, followed by overnight incubation at 4 °C with primary antibodies. After three washes with TBST, membranes were incubated with horseradish peroxidase (HRP)-conjugated secondary antibodies (1:2,000 in 5% BSA–TBST) for 1 hour at room temperature.

Signals were detected using enhanced chemiluminescence reagents, and bands were visualized with the ChemiDoc imaging system. Band intensities were quantified using ImageJ software. Primary and secondary antibodies used for Western blotting are provided in the Supplementary Information.

**Quantitative RT-PCR Analysis**

Total RNA was extracted from cultured cells or mouse brain tissues using TRIzol reagent according to the manufacturer's instructions. Complementary DNA (cDNA) was synthesized from 0.5–1 μg of total RNA using the SuperScript reverse transcription kit. Quantitative real-time PCR was performed using the CFX96™ Real-Time PCR Detection System and iQ™ SYBR Green Supermix, following the manufacturer's protocol.

Relative gene expression levels were calculated using the ΔΔCt method and normalized to housekeeping genes such as *Gapdh* or *β-actin*. Primer sequences used in this study are listed in Supplementary Information.

**AAV Production and Titeration**

Adeno-associated virus serotype 5 (AAV5) vectors expressing tdTomato (*GfaABC1D-tdTomato*) or Ccl3 and Ccl4 (*GfaABC1D-Ccl3, Ccl4*) under the control of the *GfaABC1D* promoter produced in HEK293T cells using a simplified chloroform extraction protocol, as previously described[4]. HEK293T cells were co-transfected with three plasmids: a helper plasmid (Addgene #112867), a Rep/Cap plasmid for AAV5 (Addgene #104964), and the transfer plasmid (pZAC2.1-GfaABC1D-tdTomato, Addgene #44332, or in-house–constructed pZAC2.1-GfaABC1D-T7-Ccl3 and pZAC2.1-GfaABC1D-Ccl4-Myc-DDK).

At 72 hours post-transfection, culture medium and cell lysates were collected, and virus was purified following a previously established protocol. Briefly, transfected cells were subjected to repeated freeze–thaw cycles to induce lysis, and the lysates were mixed with chloroform to extract viral particles. After centrifugation, the aqueous phase containing AAV was carefully collected and combined with polyethylene glycol (PEG 8000, final concentration 50%) to precipitate viral particles. The viral pellet was resuspended in sterile PBS and stored at 4 °C until use.

Viral genome titers were determined by quantitative PCR (qPCR) targeting the WPRE sequence using the following primers:

- Forward: 5′-CACCACCTGTCAGCTCCTTT-3′
- Reverse: 5′-AAGGAAGGTCCGCTGGATTG-3′

Standard curves were generated using serial dilutions of linearized AAV plasmid DNA. Final viral preparations yielded titers ranging from $1–2 \times 10^{12}$ genome copies/mL on average.

**Behavioral Tests**

All behavioral experiments were conducted during the light phase of the light–dark cycle. Prior to testing, mice were habituated to the behavioral testing room for 30 minutes to minimize stress-induced variability.

**Grip Strength Test**

Forelimb muscle strength and endurance were measured using the BIO-GS4 system. Mice were gently held by the tail and allowed to grasp a horizontal metal T-bar with their forepaws. A steady horizontal pull was applied until the animal released its grip, and the peak force exerted just before release was automatically recorded. Each mouse underwent five trials with at least 10 minutes of rest between trials. To ensure consistency and minimize outlier effects, the highest three values from the five attempts were averaged for analysis.

**Rotarod Test**

Motor coordination and balance were evaluated using a rotarod apparatus (B.S Technolab, Seongnam, South Korea). The test was conducted over three consecutive days. On days 1 and

2, mice underwent training sessions at a constant speed of 4 rpm for 60 seconds to acclimate to the apparatus. On the third day, testing was performed with the rotarod accelerating linearly from 4 to 44 rpm over a 4-minute period. Each mouse completed three trials, separated by 40-minute rest intervals. The latency to fall from the rod was automatically recorded by the system, and mice were returned to their home cages immediately after each trial.

**Treadmill Exhaustion Test**

To assess exercise capacity, mice were subjected to a treadmill exhaustion test using a treadmill system (Jeung-Do Bio & Plant, Seoul, South Korea). All mice first underwent a training phase consisting of 5 minutes of running at 5 rpm. After a 30–60-minute rest period, testing began.

For aged mice, treadmill speed was increased by 3 rpm every 3–5 minutes until exhaustion. For young mice, a more stringent protocol was applied: starting at 5 rpm, the speed was increased by 2 rpm every 2 minutes. Exhaustion was defined as the point at which mice failed to resume running after repeated gentle stimulation. The time to exhaustion was recorded.

**Open-Field Test**

The open-field test was conducted to assess locomotor activity and anxiety-like behavior. Each mouse was placed in the center of a square open-field arena, and its movement was recorded for 10 minutes using a digital camera connected to the ANY-maze tracking software. The total distance traveled (in meters), as well as the time spent in the center versus the periphery of the arena (in seconds), were automatically quantified by the software. All measurements were conducted under consistent lighting and noise-controlled conditions.

**Body Temperature Measurement**

Core body temperature was measured using a rectal thermometer. The probe was gently inserted approximately 2cm into the rectum of each mouse, and the temperature was recorded once a stable reading was obtained.

### Supplementary List

#### Supplementary List 1 Primers Used in this Study

| Genes | Species | Direction | Primer sequences (5'-3') | Application |
|---|---|---|---|---|
| *Apoe* | Mouse | Forward | ATCCGATCCCCTGCTCAGAC | qPCR |
| | | Reverse | GTGGATCCGCTGCCAAAAAT | |
| *B2m* | Mouse | Forward | CTGGGGTAAGCCTCAAGTTCT | qPCR |
| | | Reverse | TGGCCTGCTGTGTAAGTCTC | |
| *Bst2* | Mouse | Forward | CACAGGCAAACTCCTGCAAC | qPCR |
| | | Reverse | TCCTGGTTCAGCTTCGTGAC | |
| *Ccl3* | Mouse | Forward | GCAACCAAGTCTTCTCTCAGCG | qPCR |
| | | Reverse | GTCCGGTTTCTCTTAGTCAGG | |
| *Ccl4* | Mouse | Forward | CCCAGCTCTGTGCAAACCTA | qPCR |
| | | Reverse | TGGAGCAAAGACTGCTGGTC | |
| *Cd36* | Mouse | Forward | CGTTTCAACTCTCACACACATAAG | qPCR |
| | | Reverse | TGAGACTCTGAAAGGATCAGCAC | |
| *Cst7* | Mouse | Forward | AAGCGGGTAAGCAGAAGAGC | qPCR |
| | | Reverse | AGGGCTGGGAGGATACTGTT | |
| *Cstb* | Mouse | Forward | GAGGTGTGGACCTGACTACC | qPCR |
| | | Reverse | TGAGCTTGGCCTAACCCTTAC | |
| *Cstd* | Mouse | Forward | GTAACTCTGACACTGGCTCCG | qPCR |
| | | Reverse | GTTGGAGGACACAGCAGTCAA | |

| | | | | |
|---|---|---|---|---|
| *Cx3cr1* | Mouse | Forward | CTTCCCATCTGCTCAGGACCTC | qPCR |
| | | Reverse | CGCCCAAATAACAGGCCTCA | |
| *Dgat1* | Mouse | Forward | GGAATATCCCCGTGCACAA | qPCR |
| | | Reverse | CATTTGCTGCTGCCATGTC | |
| *Dgat2* | Mouse | Forward | CCGCAAAGGCTTTGTGAA | qPCR |
| | | Reverse | GGAATAAGTGGGAACCAGATCAG | |
| *Fasn* | Mouse | Forward | GGAGGTGGTGATAGCCGGTAT | qPCR |
| | | Reverse | TGGGTAATCCATAGAGCCCAG | |
| *Ifit1* | Mouse | Forward | GTTCTGCTCTGCTGAAAACCC | qPCR |
| | | Reverse | TGGTCACCATCAGCATTCTCT | |
| *Ifit3* | Mouse | Forward | AGATTTCTGAACTGCTCAGCCC | qPCR |
| | | Reverse | CAGAGATTCCCGGTTGACCTC | |
| *Ifn-γ* | Mouse | Forward | TATAGCTGCCATCGGCTGAC | qPCR |
| | | Reverse | GGCTTTCAATGACTGTGCCG | |
| *IL-1β* | Mouse | Forward | GCCCATCCTCTGTGACTCAT | qPCR |
| | | Reverse | AGGCCACAGGTATTTTGTCG | |
| *Isg20* | Mouse | Forward | CGAGGGAGAGATCACGGACT | qPCR |
| | | Reverse | CCACCAGCTTGCCTTTCAGA | |
| *Lgals3* | Mouse | Forward | CTAATCAGGTGAGCGGCACAG | qPCR |
| | | Reverse | AAGGCATCGTTAAGCGAAAAGC | |
| *Lipa* | Mouse | Forward | AAGCTCGCCTGCTTGTAGTG | qPCR |
| | | Reverse | TGGAGTTGCATCTTCCGGG | |

| | | | | |
|---|---|---|---|---|
| *Plin2* | Mouse | Forward | AACTTGCATTTGTCCCGTCG | qPCR |
| | | Reverse | CGGAGGACACAAGGTCGTAG | |
| *Pnpla2* | Mouse | Forward | GGAACCAAAGGACCTGATGACC | qPCR |
| | | Reverse | CATCAGGCAGCCACTCCAAC | |
| *P16* | Mouse | Forward | TGGTCACTGTGAGGATTCAGC | qPCR |
| | | Reverse | CGTGAACGTTGCCCATCATC | |
| *Tnf-α* | Mouse | Forward | ATGGCCTCCCTCTCATCAGT | qPCR |
| | | Reverse | TTTGCTACGACGTGGGCTAC | |
| *Trem2* | Mouse | Forward | AGCGATGGGAGCCTTGAGAG | qPCR |
| | | Reverse | CCCAGGATAGGTGGGCTTGA | |
| *Tyrobp* | Mouse | Forward | GCTGGGATTGTTCTGGGTGA | qPCR |
| | | Reverse | GACCTTGACGCTTCCACTGT | |

**Supplementary List 2 Antibodies Used in this Study**

| Antibodies | Supplier | Catalog | Source | Clone number | Application | Dilution |
|---|---|---|---|---|---|---|
| B2m | Abcam | ab75853 | rabbit monoclonal antibody | clone EP2978Y | IHC | 1:500 |
| CCl3 | Rndsystem | AF-450-NA | goat polyclonal antibody | / | IHC, WB | 1:200 |
| CD3 | Novus Biologicals | NB600-1441 | rabbit monoclonal antibody | clone SP7 | IHC | 1:200 |
| CD4 | Abcam | AB183685 | rabbit monoclonal antibody | EPR19514 | IHC | |
| CD8 | Invitrogen | 14-0808-80 | rat monoclonal antibody | 4SM15 | IHC | 1:800 |
| CD8 | Abcam | Ab4055 | rabbit polyclonal antibody | / | IHC | 1:800 |
| Galectin-3 | Santa Cruz Biotechnology | sc-23939 | rat monoclonal antibody | clone M1/87 | IHC, ICC | 1:500 |
| IBA1 | FUJIFILM Cellular Dynamics | 019-19741 | rabbit polyclonal antibody | / | IHC | 1:1000 |
| IBA1 | Synaptic systems | 234-308 | guinea pig monoclonal antibody | clone Gp311H9 | IHC | 1:1000 |
| IFITM3 | Abclonal | A13070-20 | rabbit polyclonal antibody | / | IHC | 1:500 |
| LAMP1 | Abcam | ab25630 | mouse monoclonal antibody | clone H4A3 | ICC | 1:1000 |
| NKX2.1 | Invitrogen | MA5-13961 | mouse monoclonal antibody | clone 8G7G3/1 | ICC | 1:800 |
| PLIN2 | Abcam | ab52356 | rabbit polyclonal antibody | / | IHC, ICC | 1:500 |
| pSTAT3 | Cell Signaling Technology | #9131S | rabbit polyclonal antibody | / | IHC | 1:1000 |
| p16 | Santa Cruz Biotechnology | sc-377412 | mouse monoclonal antibody | clone C-7 | IHC | 1:1000 |
| RAX | Takara | M228 | mouse polyclonal antibody | / | ICC | 1:500 |
| SOX2 | Millipore Sigma | AB5603 | rabbit polyclonal antibody | / | ICC | 1:1000 |
| T-bet | Abcam | ab307193 | rabbit monoclonal antibody | clone EPR27094-16 | IHC | 1:500 |

| | | | | | | |
|---|---|---|---|---|---|---|
| Vimentin | Millipore Sigma | AB5733 | chicken polyclonal antibody | / | IHC | 1:1000 |
| DAPI | Vector laboratories | H-1200-10 | polyclonal antibody | | IHC, ICC | |
| Alexa Fluor 488 goat anti-mouse | Invitrogen | A11001 | goat polyclonal antibody | / | IHC, ICC | 1:500 |
| Alexa Fluor 488 goat anti-rabbit | Invitrogen | A11008 | goat polyclonal antibody | / | IHC, ICC | 1:500 |
| Alexa Fluor 488 (plus) goat anti-rat | Invitrogen | A48262 | goat polyclonal antibody | / | IHC, ICC | 1:500 |
| Alexa Fluor 488 goat anti-chicken | Invitrogen | A32931 | goat polyclonal antibody | / | IHC | 1:500 |
| Cy3 goat anti-rabbit | Jackson immunoresearch | 111-165-144 | goat polyclonal antibody | / | IHC, ICC | 1:500 |
| Cy3 donkey anti-chicken | Jackson immunoresearch | 703-165-155 | donkey polyclonal antibody | / | IHC | 1:500 |
| Cy3 goat anti-mouse | Jackson immunoresearch | 115-165-146 | goat polyclonal antibody | / | IHC, ICC | 1:500 |
| Cy3 goat anti-Guinea Pig | Jackson immunoresearch | 106-165-003 | goat polyclonal antibody | / | IHC | 1:500 |
| CASP3 | Cell Signaling Technology | #14220 | rabbit monoclonal antibody | clone D3R6Y | WB | 1:1000 |
| CD3e | Abcam | ab16669 | rabbit monoclonal antibody | clone SP7 | WB | 1:400 |
| CD3e | Novus Biologicals | NB600-1441 | rabbit monoclonal antibody | clone SP7 | WB | 1:400 |
| IL-1β | Cell Signaling Technology | #12242 | mouse monoclonal antibody | clone 3A6 | WB | 1:1000 |
| pSTAT1 | Cell Signaling Technology | #9167S | rabbit monoclonal antibody | clone 58D6 | WB | 1:1000 |
| STAT1 | Cell Signaling Technology | #14994T | rabbit monoclonal antibody | clone D1K9Y | WB | 1:1000 |
| TNF-α | Abcam | ab307164 | rabbit multiclonal antibody | clone RM1005 | WB | 1:1000 |
| β-actin | Invitrogen | MA1-140 | mouse monoclonal antibody | clone 15G5A11/E2 | WB | 1:5000 |
| mouse IgG, HRP-linked Antibody | cell signaling | 7076S | polyclonal antibody | | WB | 1:2000 |
| rabbit IgG, HRP-linked Antibody | cell signaling | 7074S | polyclonal antibody | | WB | 1:2000 |
| HRP-conjugated donkey anti-goat IgG | Jackson immunoresearch | 705-035-147 | donkey polyclonal antibody | / | WB | 1:2000 |

**Supplementary List 3 Reagents Used in this Study**

| Reagent or Resource | Source | Catalog |
|---|---|---|
| Poly-L-Ornithine | Sigma-Aldrich | P3655 |
| Fibronectin | Sigma-Aldrich | F1141 |
| modified N2 medium | Thermo Fisher Scientific | 12500-062 |
| DMEM/F12 medium | Thermo Fisher Scientific | 11320-033 |
| DMEM | Thermo Fisher Scientific | 11995-065 |
| RPMI-1640 | Corning | 10-040-CV |
| Recombinant human insulin | Gibco | 12585-014 |
| Recombinant human FGF basic | R&D Systems | 233-FB |
| B-27 supplement without vitamin A | Thermo Fisher Scientific | 12587010 |
| B-27 supplement | Gibco | 17504044 |
| GlutaMAX | Gibco | 35050061 |
| ACK buffer | Thermo Fisher Scientific | A1049201 |
| Accutase | Stem cell | #07922 |
| Penicillin-streptomycin | Thermo Fisher Scientific | 15140122 |
| FBS | Thermo Fisher Scientific | A5670701 |
| Trypsin-EDTA | Thermo Fisher Scientific | 25300054 |
| D-glucose | Sigma Aldrich | G7021 |
| Horse serum | | |
| Hank's Balanced Salt Solution | Gibco | 14065-056 |
| Phosphate-buffered saline | Gibco | 70011-044 |
| Sucrose | JUNSEI | 31365-0350 |
| Citrate buffer (pH 6.0) | Sigma Aldrich | C9999 |
| Triton X-100 | Sigma Aldrich | X100 |
| oxidized low-density lipoprotein | Invitrogen | L34357 |
| ceramide | Tocris bioscience | No. 0744 |
| palmitic acid | Sigma | P0500 |
| cholesterol | Sigma | C4951 |
| Oleic acid | Sigma | O1383 |
| low-density lipoprotein | Invitrogen | L3486 |
| Dynabeads™ Mouse T-Activator CD3/CD28 | Gibco | 11456D |
| maraviroc | Selleckchem | S2003 |
| cenicriviroc | MedChemExpress | HY-14882 |
| 4% paraformaldehyde | Wako Chemicals | 163-20145 |
| TRIzol reagent | Invitrogen | 15596018 |
| Avidin/Biotin Blocking Kit | Vector Laboratories | SP-2001 |
| ABC reagent and streptavidin–HRP | Vector Laboratories | PK-6100 |
| DAB Substrate Kit | Vector Laboratories | SK-4100 |
| TrueBlack | Biotium | 23014 |

| | | |
|---|---|---|
| Annexin V/PI Apoptosis Detection Kit | Invitrogen | V13242 |
| Pierce BCA Protein Assay Kit | Thermo Fisher Scientific | 23225 |
| enhanced chemiluminescence reagents | Thermo Fisher Scientific | 34580 |
| protease inhibitor cocktail | Roche | 11873580001 |
| phosphatase inhibitor cocktail | Sigma-Aldrich | 4906845001 |
| EthD-1 | Invitrogen | E1169 |
| Oxidized LDL ELISA Kit | CUSABIO | CSB-E07933 |
| Mouse MIP-1 alpha (CCL3) ELISA kit | Invitrogen | 88-56013-22 |
| Oil Red O stain kit (Lipid stain) | Abcam | AB150678 |
| Quant-iT RiboGreen RNA assay kit | Invitrogen | R11490 |
| TruSeq Stranded mRNA Sample Prep Kit | Invitrogen | 20020595 |
| KAPA Library Quantification Kit for Illumina | KAPA Biosystems | KK4854 |
| SuperScript reverse transcription kit | Thermo Fisher Scientific | 18080044 |
| iQ™ SYBR Green Supermix | Bio-rad | #1708882 |

**Supplementary List 4 Apparatus and Equipment Used in this Study**

| Equipment | Source | Model / Details |
|---|---|---|
| Transwell plates, 5.0 µm pore | Corning | CLS3422 |
| Multi tissue dissociation kit 1 | Miltenyi Biotec | 130-110-203 |
| gentleMACS Dissociator | Miltenyi Biotec | |
| Debris Removal Beads | Miltenyi Biotec | 130-096-433 |
| MACS Separation System | Miltenyi Biotec | 130-104-075 |
| Chromium Next GEM Single Cell 5′ & V(D)J Library Kits | 10x Genomics | |
| TapeStation RNA ScreenTape system | Agilent | #5067-5576 |
| TapeStation D1000 ScreenTape | Agilent | #5067-5582 |
| Stellaris 5 LIA confocal microscope | Leica | |
| DM5000B brightfield microscope | Leica | |
| AxioScan.Z1 slide scanner | Zeiss | |
| FACS Canto II cytometer | BD Biosciences | |
| CFX96™ Real-Time PCR System | Bio-Rad | |
| ChemiDoc Imaging System | Bio-Rad | |
| NMR body composition analyzer | EchoMRI | EchoMRI-700 |
| Metabolic cage system | TSE Systems | PhenoMaster 6026 |
| Rotarod apparatus | B.S Technolab | |
| Grip strength meter | BIOSEB | BIO-GS4 |
| Treadmill system | Jeung-Do Bio & Plant | |
| Rectal thermometer | Testo | Testo 925 |
| Stereotaxic frame | Stoelting | |
| Osmotic pumps | Alzet | Model #2002 |
| Brain Infusion Kit #3 | Alzet | |
| Surgical adhesive | Henkel | Loctite 454 |
| Sutures | Ailee | SK 434, 18 mm |
| Osmotic pump | Alzet | Model #2002 |
| Brain Infusion Kit #3 | Alzet | |

### Supplementary List 5 Software and Packages Used in this Study

| Software & Packages | Source | Website | Reference |
|---|---|---|---|
| Cell Ranger (v. 5.0.1) | 10x Genomics | https://www.10xgenomics.com/ | - |
| Cellbender (v0.3.0) | Fleming, et al., 2023 | https://github.com/broadinstitute/CellBender | 5 |
| R (v. 4.4.2) | R Core Team | https://www.r-project.org/ | - |
| Seurat (v. 5.3.0) | Hao et al., 2023 | https://satijalab.org/seurat/ | 6 |
| scDblFinder (v. 1.18.0) | Germain et al., 2021 | https://github.com/plger/scDblFinder | 7 |
| sc-type | Ianevski et al., 2022 | https://github.com/IanevskiAleksandr/sc-type | 8 |
| clusterProfiler (v. 4.12.1) | Wu et al., 2021 | https://guangchuangyu.github.io/software/clusterProfiler/ | 9 |
| Org.Mm.eg.db (v.3.19.1) | Biocore Data Team | https://bioconductor.org/packages/release/data/annotation/html/org.Mm.eg.db.html | |
| GOSemSim (v2.30) | Yu, 2020 | https://github.com/YuLab-SMU/GOSemSim | 10 |
| rrvgo (v1.16) | Sayols, 2023 | https://www.bioconductor.org/packages/release/bioc/html/rrvgo.html | 11 |
| ggplot2 (v3.5.2) | Wickham et al., 2016 | https://ggplot2.tidyverse.org | 12 |
| ggpubr (v0.6.0) | Kassambara, 2025 | https://rpkgs.datanovia.com/ggpubr/. | - |
| ggsci (v3.2.0) | Xiao et al., 2025 | https://nanx.me/ggsci/. | - |
| viridis (v0.6.5) | Garnier et al., 2023 | https://sjmgarnier.github.io/viridis/ | - |
| scCustomize (v3.0.1) | Marsh, 2024 | https://github.com/samuel-marsh/scCustomize | - |
| Nebulosa (v1.14.0) | Powell and Alquicira-Hernandez, 2021 | https://www.bioconductor.org/packages/release/bioc/html/Nebulosa.html | 13 |
| scRepertoire (v2.0.7) | Yang et al., 2024 | https://github.com/BorchLab/scRepertoire | 14 |
| ProjecTILs (v3.6.0) | Andreatta et al., 2021 | https://github.com/carmonalab/ProjecTILs | 15 |
| Fiji | Schindelin et al., 2012 | https://imagej.net/ | 16 |
